# Intracellular Galectin-9 controls dendritic cell function by maintaining plasma membrane rigidity

**DOI:** 10.1101/739797

**Authors:** Laia Querol Cano, Oya Tagit, Anne van Duffelen, Shannon Dieltjes, Sonja I. Buschow, Toshiro Niki, Mitsuomi Hirashima, Ben Joosten, Koen van den Dries, Alessandra Cambi, Carl G. Figdor, Annemiek B. van Spriel

## Abstract

Extracellular Galectins constitute a novel mechanism of membrane protein organisation at the cell surface. Although Galectins are also highly expressed intracellularly, their cytosolic functions are poorly understood. Here, we investigated the role of Galectin-9 in dendritic cell (DC) surface organisation and function. By combining functional, super-resolution and atomic force microscopy experiments to analyse membrane stiffness, we identified intracellular Galectin-9 to be indispensable for plasma membrane integrity and structure in DCs. Galectin-9 knockdown studies revealed intracellular Galectin-9 to directly control cortical membrane structure via modulating Rac1 activity, providing the underlying mechanism of Galectin-9-dependent actin cytoskeleton organisation. Consequent to its role in maintaining plasma membrane structure, phagocytosis studies revealed that Galectin-9 was essential for C-type lectin receptor-mediated pathogen uptake by human DCs. This was confirmed by the impaired phagocytic capacity of Galectin-9-null murine DCs. Together, this study demonstrates a novel role for intracellular Galectin-9 in modulating DC function, which may be evolutionary conserved.

## Introduction

<u>D</u>endritic <u>c</u>ells (DCs) constitute the major group of antigen presenting cells that constantly patrol the body for microbes and are essential for linking innate and adaptive immune responses (Banchereau and Steinman, 1998). DCs are equipped with a diverse membrane receptor repertoire to take up pathogens including Toll-like receptors, scavenger receptors and C-type lectins, such as the mannose receptor and the <u>d</u>endritic <u>c</u>ell–<u>s</u>pecific intercellular adhesion molecule <u>g</u>rabbing <u>n</u>on-integrin receptor (DC-SIGN) (Banchereau et al., 2000; Buschow et al., 2012; Geijtenbeek et al., 2000; Heinsbroek et al., 2008; Savina and Amigorena, 2007). Engagement of these receptors with their ligand is accompanied by cytoskeletal changes, which allow for the capture and engulfment of phagocytic targets (Baranov et al., 2016; Sano et al., 2003). Actin polymerisation is instrumental in forming a nascent phagosome for pathogen engulfment and actin-driven mechanical forces enable pathogen internalisation (May et al., 2000). In addition, phagocytosis is dependent on plasma membrane organisation and loss of membrane structure results in impaired pathogen recognition, defective migration and compromised immunological synapse formation (Alvarez et al., 2008; Buschow et al., 2012; Heuze et al., 2013). Galectins, a family of ß-galactoside-binding proteins, have been recently identified as a novel mechanism of membrane organisation (Elola et al., 2015; Lajoie et al., 2009; Nabi et al., 2015) due to their ability to interact with and crosslink specific carbohydrate structures. As such, Galectins can simultaneously interact with multiple glycoconjugates, thereby regulating the dynamics of glycosylated binding partners, limiting receptor internalisation and establishing membrane microdomains (Elola et al., 2015). Notably, Galectins are also abundantly expressed intracellularly although their cytosolic functions are not well characterised (Hsu et al., 2015a; Johannes et al., 2018; Liu et al., 2002; Liu and Rabinovich, 2005). Recently, Galectins have been discovered as novel regulators of several immune processes, such as T cell homeostasis, inflammation and immune disorders (de Oliveira et al., 2015; Rabinovich and Toscano, 2009; Sundblad et al., 2017; Thiemann and Baum, 2016).

Galectin-9 was first discovered as an eosinophil chemoattractant and to date, most studies have focused on studying Galectin-9 in inflammation or infection processes (Curciarello et al., 2014; Hsu et al., 2015b; Jost et al., 2013). Extracellular Galectin-9 has been implicated in inhibiting T cell immunity by promoting T cell apoptosis and differentiation into regulatory T cells (Anderson et al., 2007; Bi et al., 2008; Zhu et al., 2005). Furthermore, extracellular Galectin-9 acts as a suppressor of B cell signalling *via* binding to the B cell receptor (Cao et al., 2018; Giovannone et al., 2018). While these studies indicate that Galectin-9 plays an inhibitory role on lymphocytes, its function in myeloid cells remains poorly understood. Moreover, Galectin-9 is also highly expressed intracellularly, and although implicated in protein-protein interactions and mRNA splicing (Heusschen et al., 2013; Liu et al., 2002; Sundblad et al., 2017), the function of cytosolic Galectin-9 in the immune system continues to be ill defined.

Here, we demonstrate that intracellular Galectin-9 is essential for sustaining cortical actin cytoskeleton rigidity and phagocytosis in DCs. Our work indicates a novel evolutionary conserved mechanism by which intracellular Galectins stabilise plasma membrane structure by actin cytoskeleton reorganisation.

## Results

### Galectins are essential in governing human dendritic cell function

The role of Galectins in the initiation of the immune response is poorly understood and although Galectin-3 has been implicated in macrophage-mediated uptake in mice (Sano et al., 2003), no studies have been performed to elucidate Galectin function in dendritic cells (DCs). To address this question, we generated DCs lacking Galectin-3 and/or Galectin-9 by electroporating human <u>mo</u>nocyte-derived <u>d</u>endritic <u>c</u>ells (moDCs) with either a specific *galectin* siRNA (*gal3* and/or *gal9*) or a <u>n</u>on-targeting (NT) siRNA control prior to challenge them with FITC-labelled zymosan particles, a fungal cell wall extract (de la Rosa et al., 2005). Subsequent immunolabelling without permeabilisation using an antibody directed against FITC allowed for selective labelling of membrane bound particles. Galectin-9 and -3 protein knockdowns were confirmed by flow cytometry showing that both proteins were depleted to a similar extent (70 %, Fig. 1A and 1B). The efficiency of Galectin knockdown was comparable between cells transfected with a single siRNA or with dual siRNA, and Galectin-9 knockdown did not affect Galectin-3 expression or *vice versa* (Fig. S1). Depletion of Galectin-9 impaired particle uptake to a greater extend when comparing to that observed upon Galectin-3 knockdown (Fig. 1C, 1D and 1E). Moreover, there was no additive effect of knocking-down both Galectin-9 and -3 (Fig. 1E). Taken together, this data demonstrates that Galectins are required for phagocytosis by DCs, and indicates that Galectin-9 is a major player in this process.

**Figure 1.**
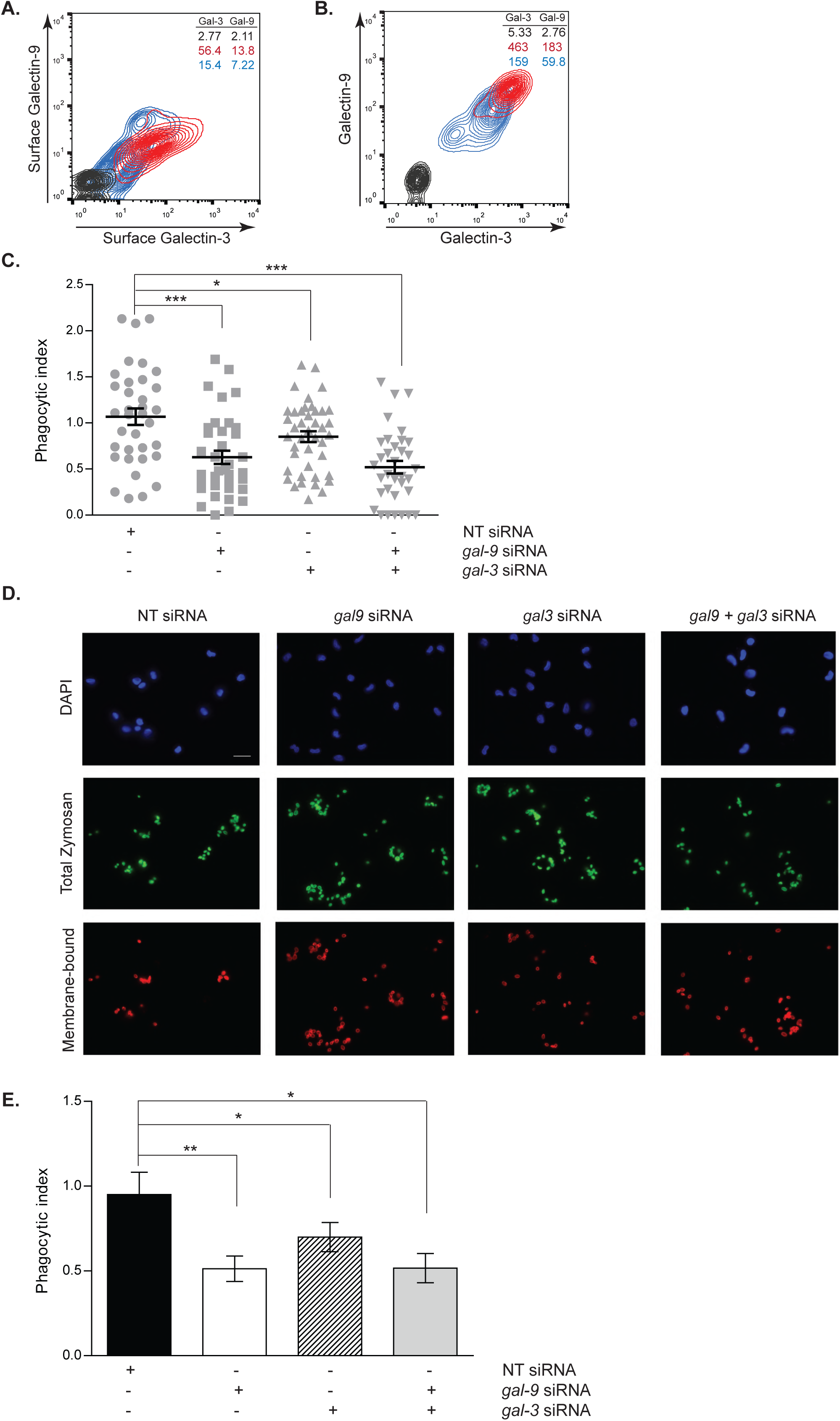
Galectin-9 is required for dendritic cell function. **A** and **B** moDCs were transfected with *gal9* siRNA and/or *gal3* siRNA or a non-targeting siRNA (NT). Surface only (A) and total (B) Galectin-9 and Galectin-3 knockdown were confirmed by flow cytometry 48 h after transfection. Red population = NT siRNA; blue population = *gal9* and *gal3* siRNA transfected moDCs; black population = isotype control. Numbers in inset indicate geometrical mean fluorescence intensity (gMFI). **C**. NT, *gal9* and/or *gal3* siRNA-transfected cells were challenged with zymosan for 60 min, after which cells were fixed, stained and the phagocytic index calculated. Graphs show representative results for one donor. Each dot represents phagocytic index obtained for one image field. 20-30 image fields were analysed per condition and each image field contained 10-20 cells. **D**. Representative images from results shown in (C). **E.** Quantification and statistical analysis of experiments depicted in (D). Results show the mean value ± SEM for four independent donors. Unpaired students t-test was conducted between NT and *gal9* siRNA and between NT and *gal3* siRNA-transfected cells. * p < 0.05, ** p < 0.005, *** p < 0.0001.

### Galectin-9 is essential for phagocytosis by dendritic cells

We previously identified Galectin-9 as part of the DC-SIGN, a phagocytic receptor present in immature DCs, -mediated phagosomes, although no functional studies were performed to assess the role of Galectin-9 in DC function (Buschow et al., 2012; Cambi et al., 2003; Geijtenbeek et al., 2000; Liu et al., 2017; Manzo et al., 2012). Co-immunoprecipitation experiments revealed DC-SIGN association with Galectin-9 in DCs, demonstrating their molecular interaction (Fig. 2A). To investigate the role of Galectin-9 in DC-SIGN-mediated phagocytosis, Galectin-9 knockdown (Gal-9 KD) and NT control (referred to as wild type (WT)) DCs were challenged with zymosan particles. Galectin-9 protein knockdown (90 %) was confirmed by flow cytometry (Fig. S2A) and Western Blotting (Fig. S2B). No significant differences in zymosan binding were observed between NT and *gal9* siRNA-transfected DCs (Fig. 2B) implying Galectin-9 is not required for particle binding. To study the involvement of Galectin-9 in particle uptake, the phagocytic index was calculated for each of the conditions and specified time points (Fig. 2C, 2D and 2E). Galectin-9 knockdown resulted in impaired zymosan internalisation 60 min after challenging moDCs but not in earlier time points, indicating that Galectin-9 is involved in late stage zymosan uptake (Fig. 2E). Galectin-9 knockdown did not alter DC-SIGN membrane expression or receptor internalisation excluding that the uptake defect was due to deficient receptor surface levels (Fig. S3 and S4). Zymosan uptake experiments were also performed with murine <u>b</u>one <u>m</u>arrow-derived <u>d</u>endritic <u>c</u>ells (BMDCs) from wild type and galectin-9-deficient (*galectin9* ^-/-^) mice. In line with human DCs, lack of Galectin-9 in murine DCs resulted in defective phagocytic capacity, suggesting an evolutionary conserved role for Galectin-9 in phagocytosis (Fig. 3).

**Figure 2.**
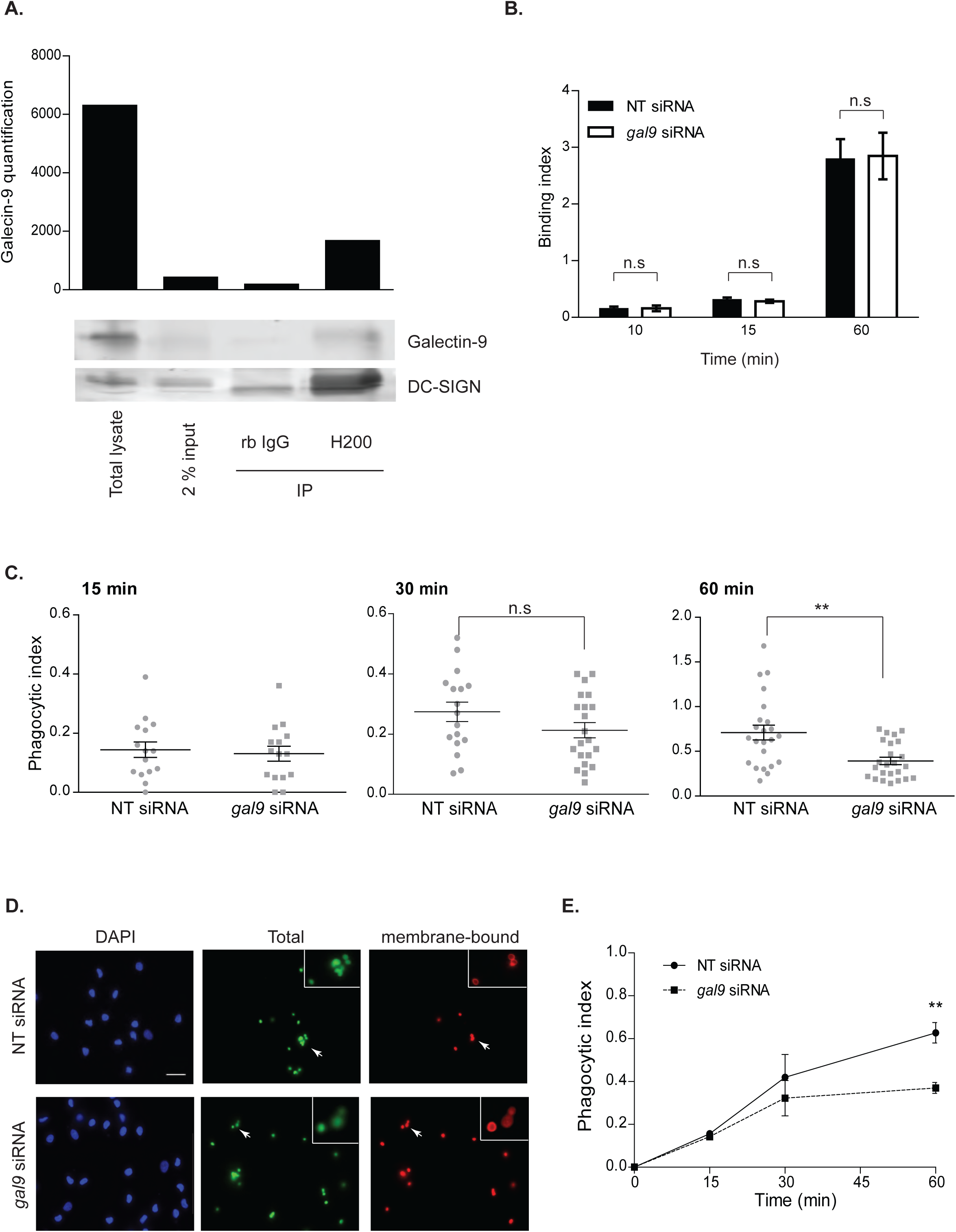
Galectin-9 is required for optimal phagocytic capacity in DCs. **A**. moDCs were lysed and whole-cell extract prepared for incubation with anti-DC-SIGN antibody (H200) or isotype control (total rabbit IgG). Immunoprecipitated (IP) complexes were resolved and probed with DC-SIGN and Galectin-9 specific antibodies. Graph shows quantification of Galectin-9 content of each sample using ImageJ. **B.** NT or *gal9* siRNA-transfected cells were challenged with zymosan for the indicated time points. After this time, cells were fixed, stained and binding index calculated for each frame analysed. Data represents mean average binding index ± SEM of three independent donors. Twenty frames were analysed for each donor and transfection. **C.** NT or *gal9* siRNA-transfected cells were treated as per (B) and phagocytic indexes calculated for each time point. Graphs show representative results for one donor. Each dot represents phagocytic index obtained for each microscopical field, each of which contained 10-20 cells. **D.** Representative images from results shown in (C) 60 min after challenging moDCs with zymosan. Scale bar: 25 μm. **E.** Quantification and statistical analysis of experiments shown in (D). Results show the mean value ± SEM for four independent donors. Unpaired students t-test was conducted between NT and *gal9* siRNAtransfected cells for all time points. n.s p> 0.05, ** p < 0.005.

**Figure 3.**
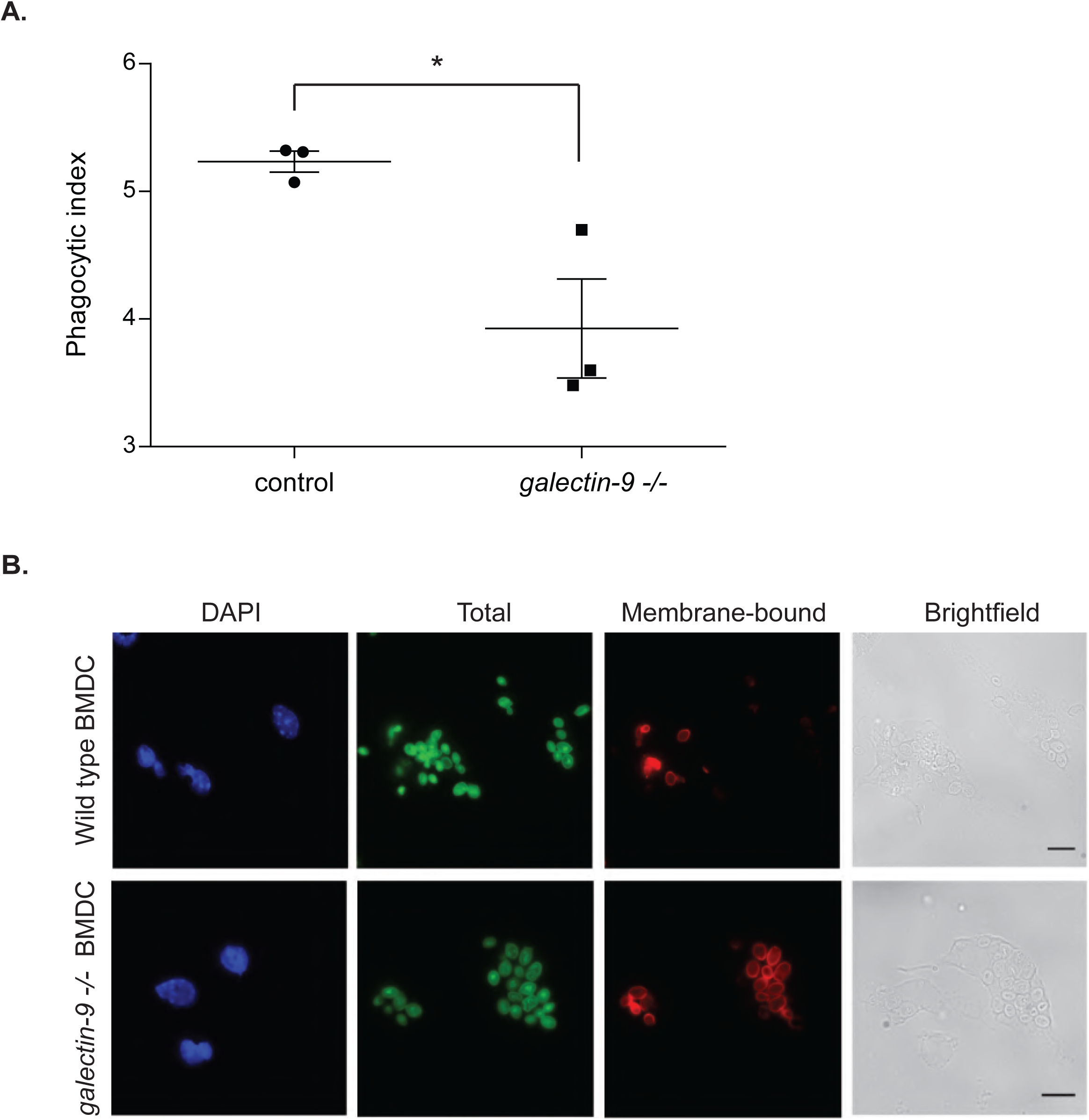
Galectin-9 function in DC is conserved between mouse and human. Bone marrow-derived dendritic cells (BMDCs) obtained from either wild type (control) or galectin9-null (galectin-9 ^-/-^) mice were seeded on coverslips, challenged with zymosan for 60 min and the phagocytic index calculated. **A.** Results show the mean phagocytic index value ± SEM for three independent mice. Unpaired students t-test was conducted between wild type and galectin-9 ^-/-^ mice. * p < 0.05. **B.** Representative images from results shown in (A). Scale bar: 10 μm.

Next, WT and Gal-9 KD moDCs were incubated with a DC-SIGN blocking antibody (clone AZN-D1) or isotype control prior to challenging them with zymosan particles. AZN-D1 does not induce DC-SIGN signalling and has a modified Fc region that cannot be recognized by the Fc receptors expressed on DCs (Geijtenbeek et al., 2000; Tacken et al., 2005). As expected, blocking DC-SIGN resulted in defective zymosan uptake by NT-transfected moDCs although zymosan uptake was unaffected by the addition of isotype controls (Fig. S5A and S5B). Analysis performed on multiple donors confirmed our observations and zymosan uptake was significantly impaired upon DC-SIGN blocking indicating that DC-SIGN is the major receptor for zymosan in DCs (Fig. S5C). Furthermore, the synergism observed between Galectin-9 depletion and DC-SIGN blocking suggests that additional surface receptors involved in zymosan uptake (Sung et al., 1983; Xia et al., 1999) are also affected by Galectin-9 knockdown (Fig. S5C). These results demonstrate that Galectin-9 is an essential component in DC-SIGN receptor-mediated uptake my DCs.

### Intracellular Galectin-9 controls plasma membrane structure in dendritic cells

To examine whether the extra- or the intracellular pool of Galectin-9 was responsible for the defect in phagocytosis, moDCs were treated with lactose prior to being challenged with zymosan particles. Lactose impairs cell surface glycan-based interactions mediated by Galectins by competing for their major ligands, which dissociates Galectins from the cell surface (Cambi et al., 2009; Lajoie et al., 2007). Although lactose treatment effectively reduced the surface levels of Galectin-9 and -3 (Fig. 4A and 4B), no effects on zymosan uptake were observed compared to untreated cells (Fig. 4C). This demonstrates that the intracellular pool of Galectin-9 is responsible for particle uptake by moDCs, and that particle binding and internalisation are independent of Galectin-9-mediated interactions at the cell surface. These findings led us to hypothesise that Galectin-9 may interact with specific cytoskeleton components, which could alter the stability and/or the formation of phagosomes. To address this, Gal-9 KD and WT DCs were analysed for their uptake ability upon treatment with cytochalasin D (cytD), which blocks actin polymerisation *via* its binding to actin filaments. Strikingly, addition of cytD resulted in a significant decrease of particle uptake in WT cells in contrast to Gal-9 KD moDCs that were not affected by cytD (Fig. 5A). We excluded that this was due to a difference in particle binding between WT and Gal-9 KD cells (Laia Querol Cano, unpublished observations) or cell viability, which was not affected upon treatment with cytD (Fig. S6A). Similar results were obtained when DCs were treated with increasing concentrations of cytD, confirming *gal9* siRNA-transfected cells to be less sensitive to actin disruption than their WT counterparts (Fig. S6B). To corroborate an impairment in the actin cytoskeleton upon Galectin-9 depletion, levels of F-actin were measured in WT and Gal-9 KD cells moDCs. A decrease of approximately 20 % in the total levels of F-actin was seen in moDCs depleted for Galectin-9 confirming a specific effect for this lectin in the actin cytoskeleton arrangement (Fig. 5B and 5C). Experiments performed with *galectin-9* ^-/-^ murine BMDCs confirmed this defect in cellular actin content (Fig. 6). Furthermore, confocal imaging showed that the percentage of F-actin positive phagosomes was reduced upon Galectin-9 knockdown in moDCs challenged with zymosan particles by approximately 40 % (Fig. 5D and 5E) in line with our previous observation (Fig. 5B). This data suggests that depletion of Galectin-9 leads to reduced actin filament formation both under basal conditions and around phagosomes. To further verify Galectin-9 involvement in directly controlling the actin cytoskeleton, super-resolution laser scanning microscopy was performed on ventral plasma membrane sheets of moDCs. These studies demonstrated that Galectin-9 closely associates with the cortical actin cytoskeleton under basal conditions (Fig. 7).

**Figure 4.**
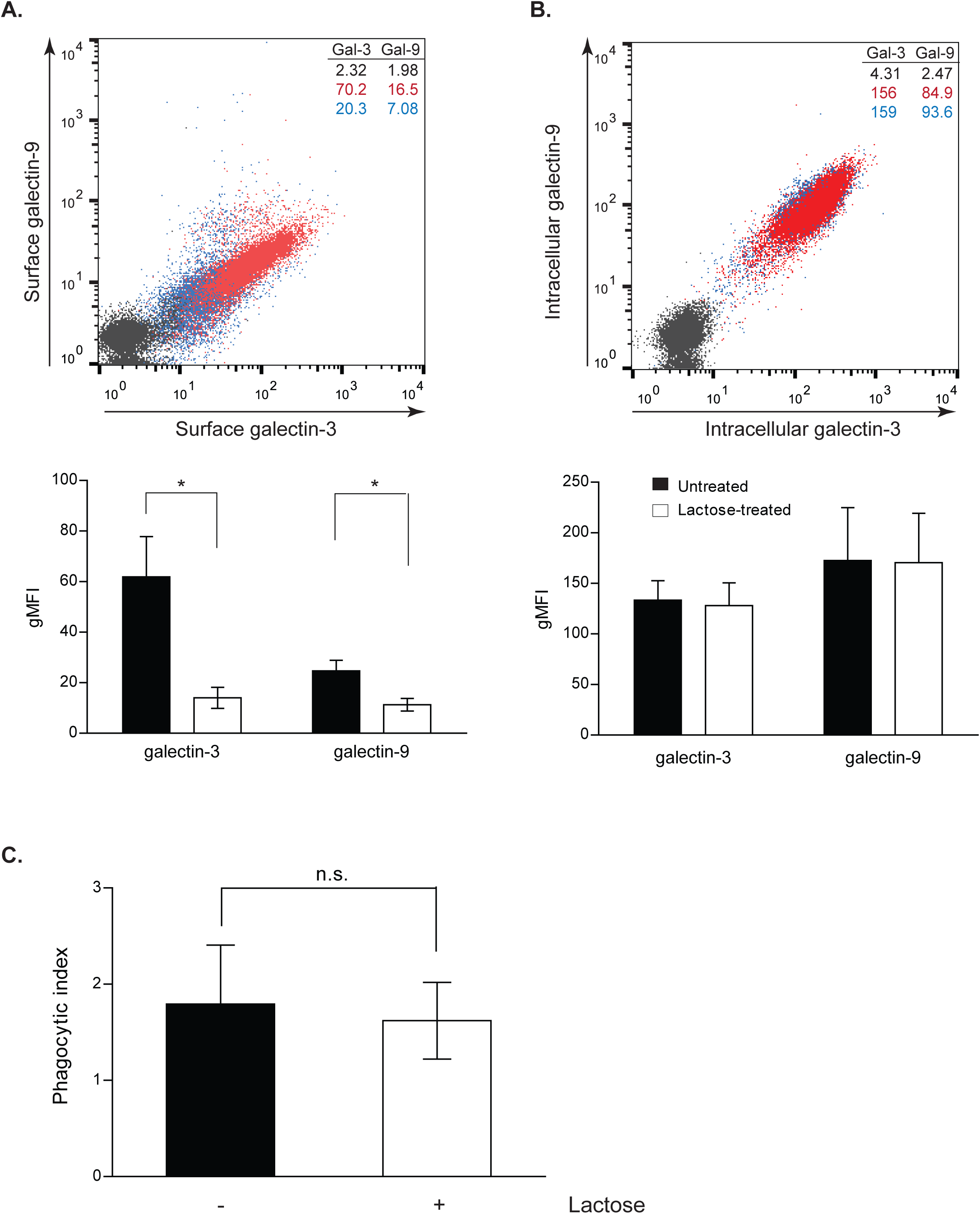
Intracellular Galectin-9 is responsible for modulating dendritic cell function. moDCs were treated with 35 mM lactose for 48 h and removal of extracellular (**A**) but not intracellular (**B**) Galectin-9 and Galectin-3 expression was confirmed by flow cytometry. Red population = untreated cells; blue population = lactose treated cells; black population = isotype control. Numbers in inset indicate gMFI. Panels depict representative results for one donor and graphs show the mean phagocytic index ± SEM for three independent donors. Unpaired students t-test was conducted between untreated control and lactose-treated cells. **C.** Control or lactose-treated cells were challenged with zymosan for 60 min. Cells were then fixed, stained and the phagocytic index calculated. Graphs show the mean value ± SEM for three independent donors. Unpaired students t-test was conducted between untreated control and lactose-treated cells. n.s p > 0.05, * p < 0.05.

**Figure 5.**
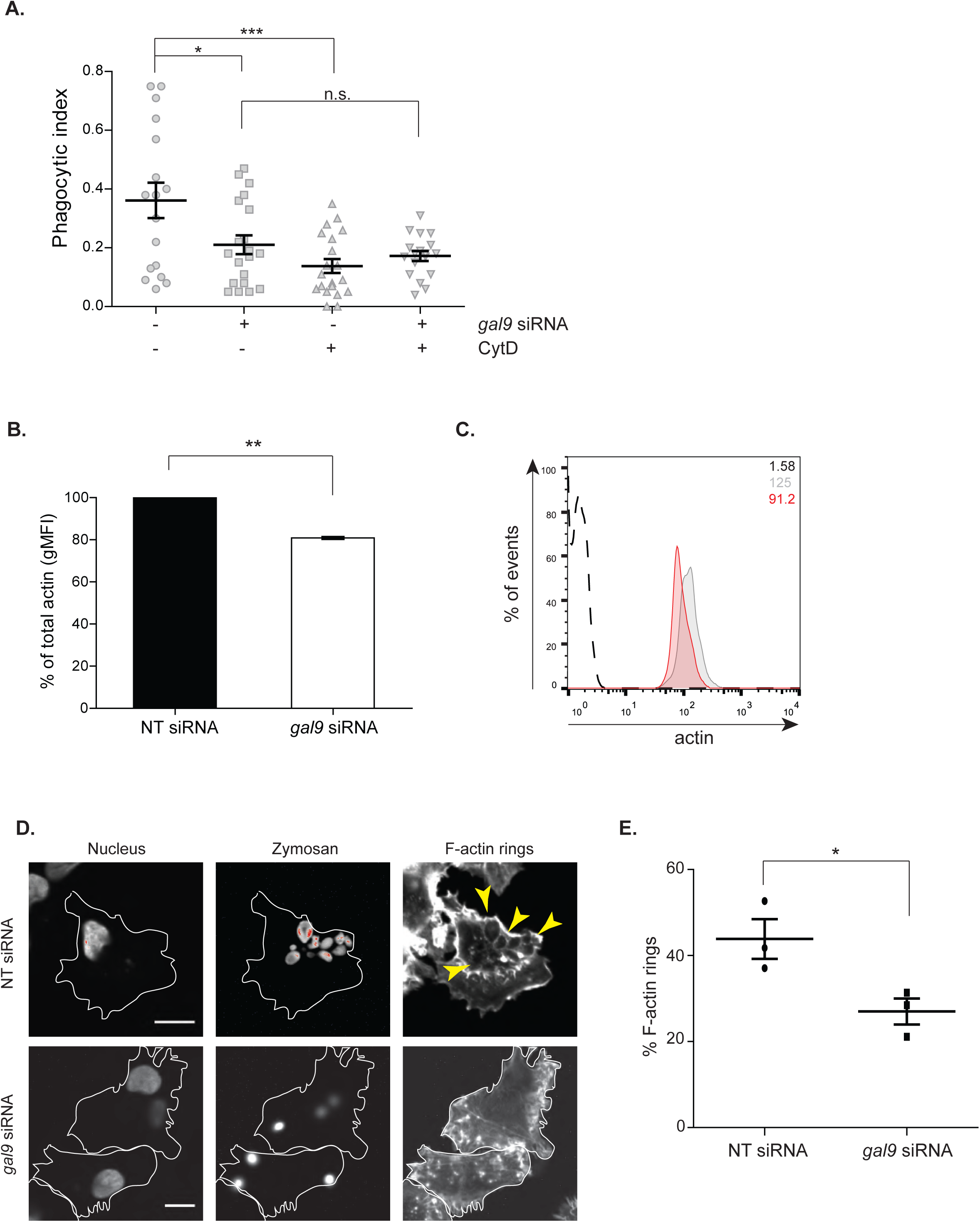
Galectin-9 modulates cellular actin cytoskeleton. **A**. NT or *gal9* siRNA-transfected moDCs were pretreated with 1.25 μg/ml cytochalasinD (cytD) for 10 min prior to being challenged with zymosan for 60 min. Cells were then fixed, stained and the phagocytic index calculated. Twenty frames were analysed for each condition and donor. Data represents mean average phagocytic index ± SEM for one representative donor out of three independent experiments. Unpaired students t-test were conducted between NT and *gal9* siRNA-transfected cells. **B.** Actin levels were analysed by flow cytometry in NT and *gal9* siRNA-transfected moDCs. Results are expressed as percent gMFI of *gal9* siRNA-transfected cells relative to their NT control. Data represents mean average % gMFI for three independent experiments ± SEM. One-way t-test was conducted. **C.** Representative histogram for actin expression after Galectin-9 knockdown. Grey area: NT siRNA-transfected moDCs; red area: *gal9* siRNA-transfected moDCs; black dotted line: isotype control. Numbers in inset indicate gMFI. **D**. Representative confocal images of F-actin rings in moDCs transfected as in (A) and challenged with zymosan particles for 15 min. **E**. Quantification of the percentage of F-actin positive phagosomes of experiments shown in (D). Data represents mean average ± SEM. n.s p> 0.05, * p < 0.05, ** p < 0.005, *** p < 0.001.

**Figure 6.**
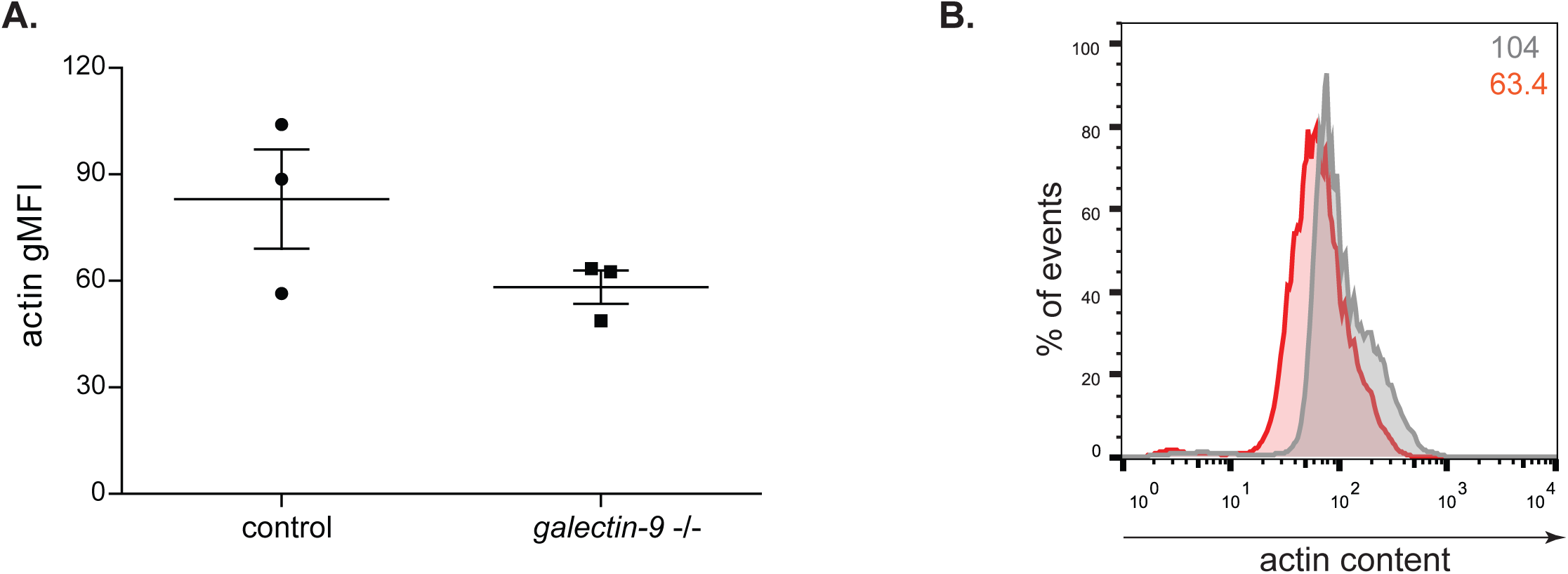
galectin-9 ^-/-^ murine DC have lower actin content. **A.** Actin levels of BMDC obtained from wild type (control) or galectin-9 ^-/-^ mice were analysed by flow cytometry. Data represents mean average gMFI for three independent mice ± SEM. **B.** Representative histogram for actin expression. Grey line: control BMDCs, red line: galectin-9 ^-/-^ BMDCs. Numbers in inset indicate gMFI.

**Figure 7.**
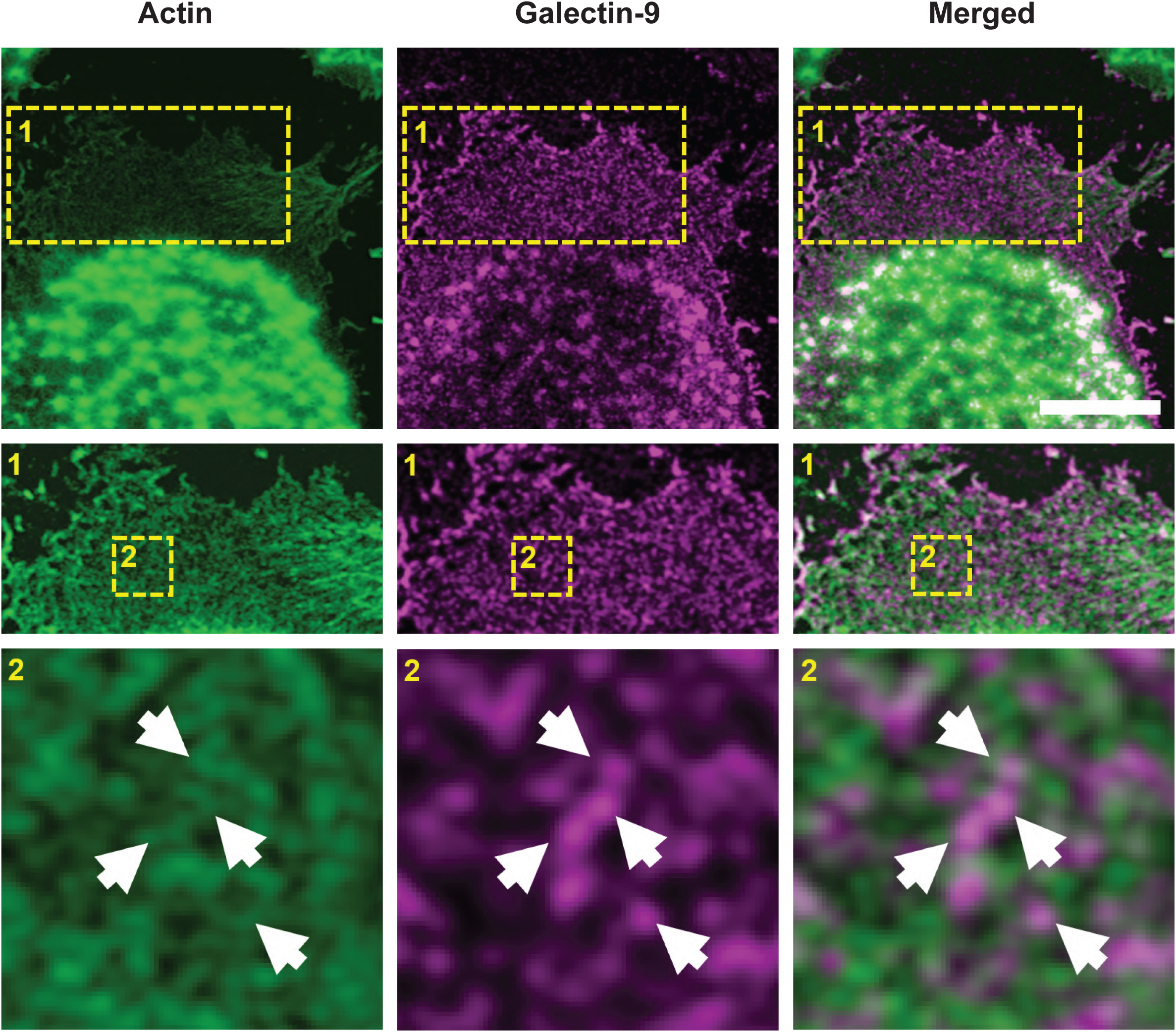
Galectin-9 closely associates with the actin cytoskeleton. Ventral plasma membrane sheets from day 5 moDCs were stained for actin and Galectin-9 and imaged with super-resolution microscopy. A representative plasma membrane sheet out of four independent experiments is shown. Gamma correction (0.2) was applied to enhance the contrast of the actin image. Lower images: magnification of the area indicated in the upper images. Scale bar:10 µm. Arrows indicate sites of Galectin-9 and actin colocalisation.

To unravel the mechanism underlying Galectin-9 function in plasma membrane integrity and structure, we exploited <u>a</u>tomic force <u>m</u>icroscopy (AFM) to analyse the cellular stiffness of moDCs transfected with either NT or *gal9* siRNA. Nanomechanical probing of the cells was achieved by obtaining a series of force-distance curves on selected points of the cell surface. A sharp non-functionalised cantilever with a radius of approximately 35 nm was brought into contact with a flat area of a single DC attached to a glass coverslip applying a mechanical force (Fig. 8A). The use of a combined brightfield-AFM setup allowed for accurately positioning the cantilever over specific areas of interest on the cell surface (Fig. 8B). Analysis of the approach force-distance curves obtained for each point of interest allowed calculation of the cytoskeletal stiffness using the linearised Sneddon equation (Fig. 8C). For this purpose, minimum and maximum fit boundaries (shown in blue) were defined respectively as 10 % and 70 % of maximum force after baseline correction (Fig. 8C). The portion of the curves that was used for fitting with the linearised model is shown in purple and the separation distance that corresponds to this fit region is in de range of 200-600 nm (Fig. 8C). Given that the thickness of the lipid bilayer is approximately 4 nm (Yokokawa et al., 2008), it is plausible to presume that the underlying cytoskeletal structures such as cortical actin and peripheral cytoplasm were also probed in our experiments. The mechanical characterisation performed on DCs shows that moDCs lacking Galectin-9 have a decreased cytoskeletal rigidity compared to their WT counterparts (Fig. 8D), in line with the defect in their cytoskeleton previously observed.

**Figure 8.**
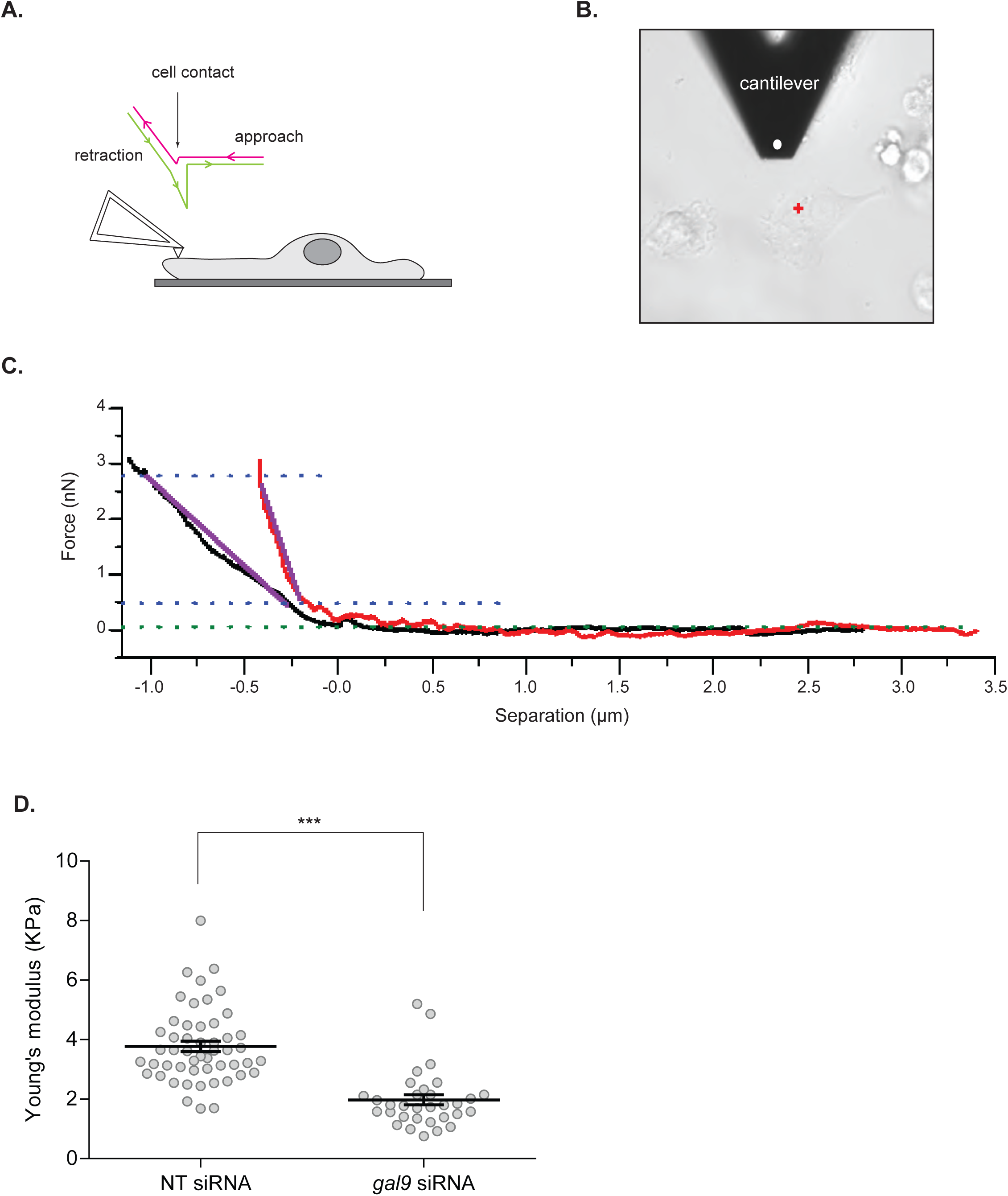
Galectin-9 alters cytoskeletal membrane rigidity. **A.** Schematic AFM single-cell elasticity measurement setup showing an overview of the AFM cantilever in contact with the DC and the cantilever movement used to measure the change in cantilever deflection. **B.** Optical image of *gal9* siRNA transfected moDCs obtained during the mechanical probing of the cell. White dot shows the position of the tip and the red cross depicts the region of interest at the membrane that was indented with the cantilever. **C.** Representative force-distance curves obtained on *gal9* siRNA transfected (black) and NT siRNA (red) moDCs and curve fitting approach to determine the Young’s modulus of elasticity. Blue lines correspond to upper and lower fit boundaries (70 % and 10 % of the maximum force respectively); purple lines show fitted portion of the curves used to calculate Young’s modulus. **D.** Young’s modulus of elasticity was calculated by fitting the force-distance curves indicated in (C). Data represents mean average Young’s modulus of elasticity ± SEM of three independent donors and each data point shows the average value for three different locations for each moDC. Ten to thirty cells were analysed for each donor in each independent experiment. Unpaired students t-test was conducted between NT and *gal9* siRNA-transfected cells.

Since it is well known that actin polymerisation is mediated by small GTPases of the Rho family including Rac1 (Caron and Hall, 1998; May et al., 2000; Norman et al., 1996; Swanson, 2008), we investigated the effect of Galectin-9 depletion on Rac1 activation by specifically measuring its GTP-bound fraction using a G-LISA colorimetric assay. Incubation of control moDCs with zymosan particles resulted in a fast induction of Rac1-GTP activity already after 5 min stimulation (Fig. 9A and 9B), which was sustained in time (Fig. 9A and 9C). Depletion of Galectin-9 abrogated Rac1 induction and no increase in its GTP-bound form could be observed upon zymosan stimulation in moDCs transfected with *gal9* siRNA at any of the time points analysed (Fig. 9A, 9B and 9C). The recruitment of total Rac1 to nascent phagocytic cups was also impaired upon Galectin-9 knockdown (Fig 9D, 9E and 9F), suggesting that Galectin-9 promotes both Rac1 recruitment and activity on phagosomes.

Taken together, intracellular Galectin-9 controls plasma membrane structure via modulating Rac1 activity and actin polymerisation, which underlies Galectin-9 requirement for phagocytosis in DCs.

**Figure 9.**
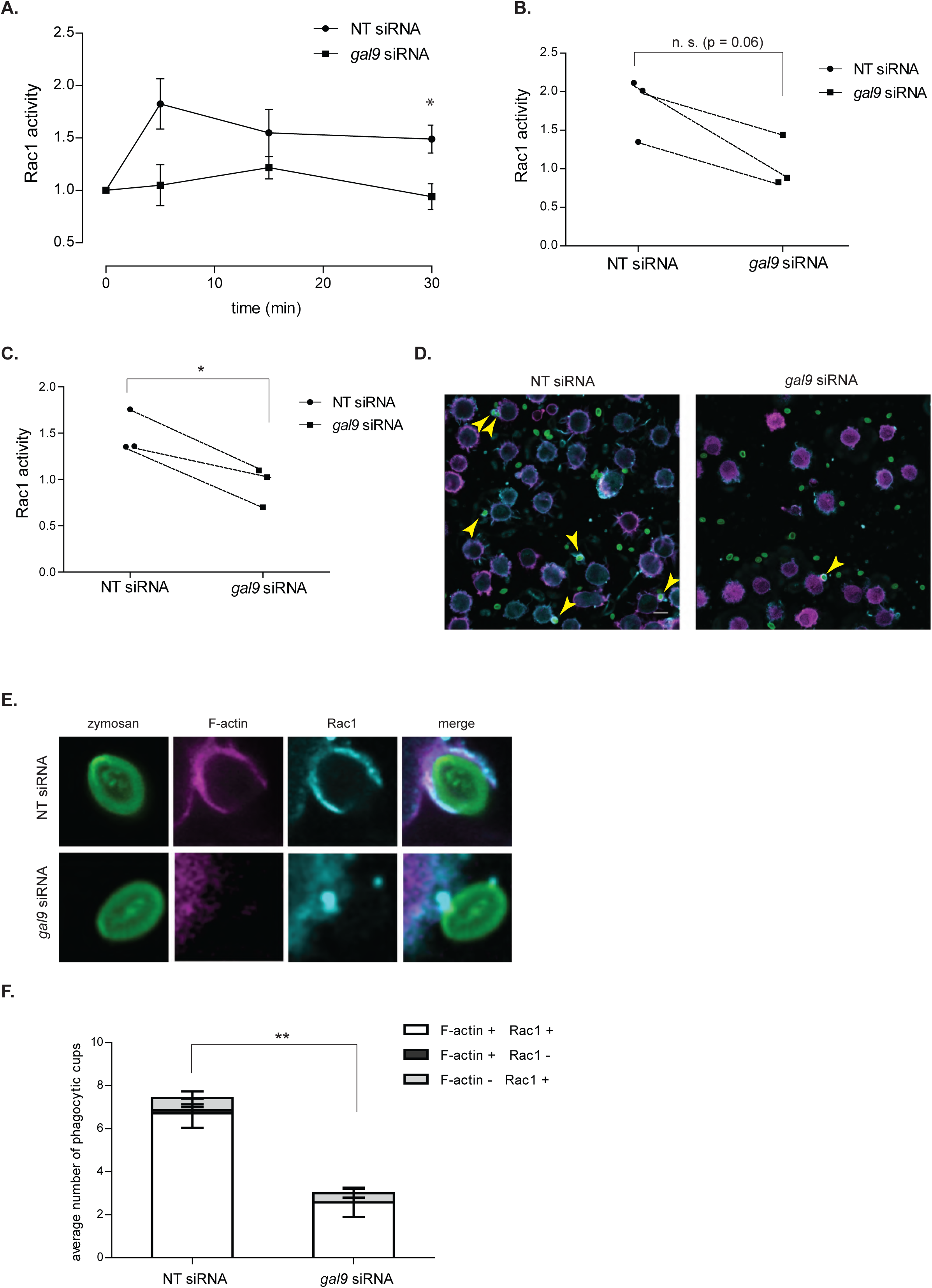
Galectin-9 promotes phagosomal Rac1 activity. **A** NT and *gal9* siRNA-transfected moDCs were challenged with zymosan particles for the indicated time points. After this time, cells were lysed, total protein was quantified and Rac1-GTP activation was determined. Results are expressed as fold increase Rac1-GTP levels and relative to the unstimulated corresponding sample. Data represents mean average Rac1-GTP fold induction ± SEM of three independent donors. Unpaired students t-test was conducted between NT and *gal9* siRNA-transfected cells. **B** and **C** each symbol represents one independent donor and lines connect paired NT and *gal9* siRNA-transfected moDCs after stimulation with zymosan for either 5 min (B) or 30 min (C). **D.** NT or *gal9* siRNA-transfected moDCs were challenged with zymosan_FITC for 5 min prior to being stained for F-actin (magenta) and Rac1 (blue) and imaged with super-resolution microscopy. A representative confocal image out of 9 images is shown. Scale bar: 10 µm. Arrows indicate overlap between Rac1 and F-actin signal on phagocytic cups. **E.** Magnification of representative phagocytic cups in NT and *gal9* siRNA-transfected moDCs treated as in (D). **F.** Number of Rac1 and/or F-actin positive phagocytic cups found on moDCs treated as in (D). Nine images containing between 20-30 cells were analysed for each condition. Data represents mean average number of phagocytic cups ± SEM. Unpaired students t-test was conducted between NT and *gal9* siRNA-transfected cells. * p < 0.05; ** P < 0.01; *** p < 0.001; n.s p > 0.05.

## Discussion

Galectins have gained increasing interest for their role as extracellular organisers of plasma membrane components via glycan-mediated interactions. Nonetheless, their mechanism of action remains poorly understood and in particular their intracellular functions are ill-defined. (Buschow et al., 2012). Here, we identified a previously unrecognised function for intracellular Galectin-9 in actin cytoskeleton reorganisation, and report a novel functional interaction between Galectin-9 and the C-type lectin receptor DC-SIGN at the plasma membrane of DCs. Several members of the Galectin family are expressed in the cytosol and some, such as Galectin-1 or 3 are predominantly intracellular proteins (Clerch et al., 1988; Hubert et al., 1995; Liu et al., 2002; Wilson et al., 1989). Very little is known regarding the localisation and function of cytoplasmic Galectin-9 although it has been implicated in protein folding and signal transduction (John and Mishra, 2016; Vasta et al., 2012). Our study now demonstrates that the large intracellular pool of Galectin-9 is responsible for the phagocytic capacity in DCs by modulating plasma membrane structure, revealing a novel function for Galectins in cytoskeleton remodelling. This was observed in both human and murine cells, which indicates Galectin-9 as an evolutionarily conserved lectin required for maintaining the cortical cytoskeleton structure and function in DCs. Our data supports a model in which Galectin-9 is essential for DC-SIGN-mediated phagocytosis, by (1) maintaining plasma membrane and cortical actin stiffness and (2) controlling cell surface receptor function (Fig. 10). We identified that the underlying mechanism involves Galectin-9-dependent activation and recruitment of Rac1-GTP upon particle incubation, which triggers actin polymerisation and the subsequent formation of phagocytic cups.

**Figure 10.**
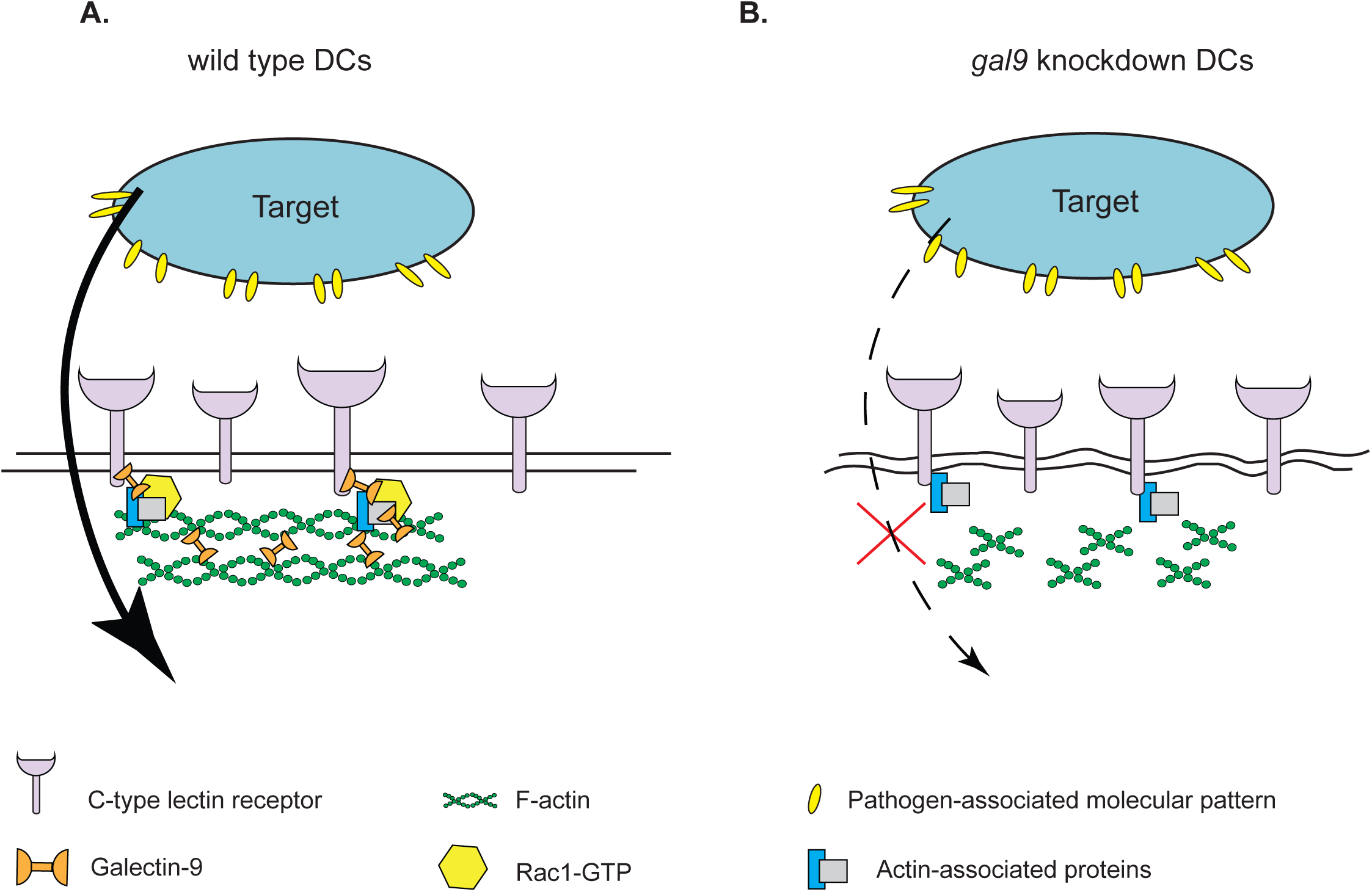
Model of the role of Galectin-9 in membrane rigidity and particle uptake. **A.** Intracellular Galectin-9 controls polymerisation of cortical actin through interacting with C-type lectin phagocytic receptors and modulating Rac1 activity, which is essential for plasma membrane integrity and successful target uptake. **B.** In the absence of Galectin-9, Rac1 activity is impaired, which results in abrogation of phagocytosis through decreased cortical actin levels and the subsequent loss of membrane rigidity.

In line with this, our AFM studies demonstrate that Galectin-9-depleted cells have a less rigid plasma membrane and cortical cytoskeleton, rendering them unable to adequately modify their structure upon particle engulfment. Moreover, inhibition of actin polymerisation did not affect the phagocytic ability of Galectin-9 knockdown cells in contrast to wild type cells. Although actin executes a pivotal function in phagocytosis, little is known regarding the mechanisms that govern F-actin recruitment to a nascent phagosome due to the lack of high-resolution data (Baranov et al., 2016). Our data supports a new concept in which intracellular Galectin-9 is required for actin polymerisation, directly controlling plasma membrane rigidity, *via* enhancing the activity of the actin-binding protein Rac1. In line with this, Galectin-1 has been recently shown to re-activate F-actin protein levels (Quinta et al., 2016). Similarly, intracellular Galectin-3 has been proven to enhance phagocytosis in macrophages *via* its interaction with F-actin in the phagocytic cups (Sano et al., 2003; Serizawa et al., 2015). Intracellular ligands have been proposed to bind Galectins through protein-protein interactions independent of carbohydrate-mediated recognition although whether both proteins interact directly or through an intermediary molecule is not known (de Oliveira et al., 2015; Johannes et al., 2018; Shimura et al., 2004). Whether Galectin-9 function in our study is carbohydrate-independent remains to be elucidated but knockdown of Galectin-9 in the presence of other Galectins was sufficient to induce defects in actin polymerisation and cellular rigidity in DCs. DC-SIGN is known to interact with actin and Lsp-1, an F-actin-interacting protein through its cytoplasmic tail (Smith et al., 2007), which likely allows for extracellular particle binding, plasma membrane deformation and actin polymerisation to occur simultaneously. Moreover, DC-SIGN signalling results in enhanced RhoA-GTPase activity (den Dunnen et al., 2009; Hodges et al., 2007). Our data now supports that depletion of intracellular Galectin-9 is sufficient to disrupt the cytosolic complex of DC-SIGN with actin-binding proteins, ultimately impairing cytoskeleton reorganisation and causing a reduction in the cellular phagocytic capacity. We identified Galectin-9 as an integral component of the actin cytoskeleton and it is conceivable that Galectin-9 exerts its effects on the cytoskeleton remodelling by directly linking F-actin filaments, actin remodelling proteins and DC-SIGN in a multi-protein complex (Fig. 9). Alternatively, Galectin-9 may connect DC-SIGN with other plasma membrane receptors known to associate with actin, such as CD44, which has been previously shown to interact with Galectin-9, as a component of DC-SIGN-directed phagosomes in DCs (Buschow et al., 2012; Wu et al., 2014).

Aside of its direct effects on DC-SIGN, we expect Galectin-9 depletion to have additional effects on the function of other lectin receptors involved in phagocytosis (mannose receptor, complement receptor 3, TLR2) (Sung et al., 1983; Xia et al., 1999). Besides CD44, Galectin-9 has been previously shown to interact with glucose transporter-2, immunoglobulin E and Tim-3, a T cell type 1 membrane protein, known to be involved in T-cell apoptosis and phagocytosis of apoptotic cells. (Wu et al., 2014; Zhu et al., 2005).

The intracellular functions of Galectin-9 and particularly its role in phagocytosis have not been previously addressed, and our work is in line with Galectin-3 and Galectin-1 function in particle uptake, highlighting the broad importance of intracellular Galectins in enhancing cellular uptake (Barrionuevo et al., 2007; Caberoy et al., 2012; Farnworth et al., 2008; Linden et al., 2013; Quattroni et al., 2012; Sano et al., 2003). Furthermore, our studies with primary DCs demonstrate that disruption of glycan interactions solely alters particle uptake but not their binding to the cell membrane, which is in agreement with previous findings (Sano et al., 2003).

In summary, our work demonstrates a novel role for intracellular Galectin-9 in the regulation of the phagocytic activity through reorganisation of the actin cytoskeleton that underlies plasma membrane rigidity in DCs. Given the plethora of cellular biological processes Galectin-9 is involved in, this novel intracellular Galectin-9 mechanism of action contributes to the general understanding of plasma membrane structure and their implications in cell function.

## Materials and methods

### Generation of monocyte-derived dendritic cells

Dendritic cells were derived from peripheral blood monocytes isolated from a buffy coat (Sanquin, Nijmegen, The Netherlands) (de Vries et al., 2002). Monocytes isolated from healthy blood donors (informed consent obtained) were cultured for up to five days in RPMI 1640 medium (Life Technologies, Bleiswijk, Netherlands) containing 10 % foetal bovine serum (FBS, Greiner Bio-one, Alphen aan den Rijn, Netherlands), 1 mM ultra-glutamine (BioWhittaker), antibiotics (100 U/ml penicillin, 100 µg/ml streptomycin and 0.25 µg/ml amphotericin B, Life Technologies), IL-4 (500 U/ml) and GM-CSF (800 U/ml) in a humidified, 5 % CO_2_. On day 3, moDCs were supplemented with new IL-4 (300 U/ml) and GM-CSF (450 U/ml).

### Generation of bone marrow-derived dendritic cells

Galectin-9-deficient mice were kindly provided by GalPharma (Takamatsu, Japan) and were described elsewhere (Seki et al., 2008). To generate bone marrow–derived DCs (BMDCs), bone marrow cells from mouse femurs were cultured in RPMI 1640 medium supplemented with 10 % FCS, 1 mM ultra-glutamine, antibiotics, and ß-mercaptoethanol in the presence of 20 ng/ml GM-CSF (PeproTech) for 8 days to generate GM-CSF BMDCs.

### Antibodies and reagents

The following primary antibodies were used for Western Blotting: rabbit anti-DC-SIGN (H200, Santa Cruz, Heidelberg, Germany) at 1:2000 (v/v), goat anti-Galectin-9 (AF2045, R&D systems, Minneapolis, Minnesota) at 1:1000 (v/v) and rat anti-tubulin (Novus Biological, Abingdon, United Kingdom) at 1:2000 (v/v). The following secondary antibodies were used: donkey anti-rabbit IRDye 680 926-32223, Li-Cor, Lincoln, Nebraska), donkey anti-goat IRDye 680 (920-32224, Li-Cor), goat anti-rat IRDye 680 (A21096, Invitrogen, Landsmeer, Netherlands). All secondary antibodies were used at 1:5000 (v/v).

The following antibodies were used for fluorescence microscopy: mouse IgG2B anti-human DC-SIGN at 2 μg/ml (DCN46, BD Biosciences, Breda, Netherlands), mouse IgG1 anti-human AZN-D1 at 2 μg/ml (Geijtenbeek et al., 2000), goat anti-human Galectin-9 at 20 μg/ml (AF23045, R&D systems); mouse monoclonal Rac1 (240106; Cell Biolabs) at 1:100, TfR (sc-65877, Santa Cruz) at 1:200, anti-FITC Alexa fluor 647 (Jackson ImmunoResearch, Huissen, Netherlands; 200-602-037) at 1:200. The following secondary antibodies were used: donkey anti-mouse IgG Alexa 647 (A31571), goat anti-mouse IgG1 Alexa 488 (A21121), donkey-anti goat IgG Alexa 488 or 647 (A11055 and A21447), rabbit anti-mouse IgG Alexa 488 (A21204) and goat anti-mouse IgG2B Alexa 647 (A21242). All secondary antibodies were purchased from Life Technologies and used at 1:400 dilution (v/v). For F-actin staining Alexa fluor-647 phalloidin (A22287, Thermofisher) or Alexa fluor-568 phalloidin (A12380, Thermofisher) were used at 1:100 dilution (v/v). To inhibit specific cytoskeleton components cytochalasin D was used (C8273, Sigma-Aldrich, Zwijndrecht, Netherlands) at a final concentration of 1.25 or 2.5 μg/ml.

### Small interfering RNA knockdown

On day 3 of DC differentiation, cells were harvested and subjected to electroporation. For Galectin-9 and Galectin-3 silencing, three custom stealth small interfering RNA (siRNA) were used. For Galectin-9 LGALS9HSS142807, LGALS9HSS142808 and LGALS9HSS142809 were used and for Galectin-3 LGALS3HSS180668, LGALS3HSS180670, LGALS3HSS180669 (Invitrogen). Equal amounts of the siRNA ON-TARGETplus non-targeting (NT) siRNA#1 (Thermo Scientific) were used as control. Cells were washed twice in PBS and once in OptiMEM without phenol red (Invitrogen). For silencing each Galectin, a total of 15 μg siRNA (5 μg from each siRNA) was transferred to a 4-mm cuvette (Bio-Rad) and 5-10×10^6^ DCs were added in 200 μl OptiMEM and incubated for 3 min before being pulsed with an exponential decay pulse at 300 V, 150 mF, in a Genepulser Xcell (Bio-Rad, Veenendaal, Netherlands), as previously described (Schuurhuis et al., 2009). Immediately after electroporation, cells were transferred to preheated (37 °C) phenol red–free RPMI 1640 culture medium supplemented with 1 % ultraglutamine, 10 % (v/v) FCS, IL-4 (300 U/ml), and GM-CSF (450 U/ml) and seeded at a final density of 5×10^5^ cells/ml.

### Co-Immunoprecipitation and Western Blotting

Endogenous DC-SIGN was immunoprecipitated from lysates of moDCs (day 6). Cells (10×10^6^) were detached using cold PBS, collected and lysed in 1 ml lysis buffer containing 150 mM NaCl, 10 mM Tris-HCl (pH 7.5), 2 mM MgCl_2_, 1 % Brij97, 2 mM CaCl_2_, 5 mM NaF, 1 mM NaVO_4_ and 1 mM PMSF for 30 min on ice. Cell lysates were pre-cleared with 3 % BSA and isotype control-coated Dynabeads (Invitrogen). Lysates were then incubated with 2 μg of anti-DC-SIGN (H200, SantaCruz) or isotype control under rotation. After incubating for 1 h at 4 °C, dynabeads were added and samples were further incubated for 2 h. Afterwards, beads were washed five times in washing buffer (150 mM NaCl, 10 mM Tris-HCl (pH 7.5), 2 mM MgCl_2_, 0.1 % Brij97, 2 mM CaCl_2_, 5 mM NaF, 1 mM NaVO_4_ and 1 mM PMSF) and bound proteins eluted in SDS sample buffer (62.5 mM Tris pH 6.8, 2 % SDS, 10 % glycerol). Proteins were separated by PAGE and blotted onto PVDF membranes. Membranes were blocked in TBS containing 3 % BSA and 1 % skim milk powder at room temperature for 1 h prior to be stained with specific antibodies against DC-SIGN and Galectin-9. Antibody signals were detected with HRP coupled secondary antibodies and developed using Odyssey CLx (Li-Cor) following manufacturer’s instructions. Images were retrieved using the Image Studio Lite 5.0 software.

### Uptake assay and immunofluorescence

2×10^5^ moDCs transfected with either non-targeting (NT) or *gal9* siRNA were seeded in phenol red free RPMI containing 10 % FBS, 1 mM ultra-glutamine, antibiotics, IL-4 (500 U/ml) and GM-CSF (800 U/ml) in a humidified, 5 % CO2-containing atmosphere for 48 hours. After this time, cells were washed with serum free RPMI and challenged with 1×10^6^ zymosan-FITC (ThermoFisher scientific) for 15, 30 or 60 min, after which moDCs were washed extensively with PBA and fixed with 4 % paraformaldehyde (PFA). When appropriate, cells were incubated with 20 μg/ml of anti-DC-SIGN (AZN-D1) or isotype control for 10 min prior to the addition of the zymosan-FITC particles. For experiments performed using cytoskeleton-blocking agents, cells were incubated with either 1.25 or 2.5 μg/ml of cytochalasin D for 10 min prior to the addition of zymosan-FITC particles. For lactose blocking experiments, moDCs were treated with growth medium containing 35 mM of lactose for 48 h prior to the addition of zymosan-FITC particles.

For immunofluorescence staining, samples were incubated without permeabilisation with an anti-FITC AF647 antibody (200-602-037, Jackson) at 1:200 (v/v) for 30 min at room temperature followed by DAPI staining (Sigma-Aldrich) at 1:3000 (v/v) for 10 min and at room temperature. Cells were then washed twice with PBS, one time with milliQ H_2_O, embedded onto glass slides using Mowiol (Calbiochem) and stored at – 20 °C until imaging. Samples were imaged with a Leica DMI6000 epi-fluorescence microscope fitted with a 63x 1.4 NA oil immersion objective, a metal halide EL6000 lamp for excitation, a DFC365FX CCD camera and GFP and DsRed filter sets (all from Leica, Wetzlar, Germany). Focus was kept stable with the adaptive focus control from Leica. Images were analysed with ImageJ software. Twenty to thirty frames were analysed for each condition and the phagocytic index ((total number of particles –total number of membrane-bound particles)/number of cells) and the binding index (number or bound particles/number of cells) calculated for each frame.

### Flow cytometry

Single cell suspensions were stained with specific antibodies or isotype control as negative control for 30 min at 4 °C. The following antibodies were used: mouse-anti DC-SIGN (clone AZN-D1, (Geijtenbeek et al., 2000)) at 5 μg/ml; goat anti-Galectin-9 (AF2045, R&D systems) at 20 μg/ml. Before staining, moDCs and U-937 cells were incubated with 2 % human serum for 10 min on ice to block non-specific interaction of the antibodies with FcRs. All secondary antibodies were conjugated to Alexa Fluor 488 or Alexa Fluor 647 dyes (Invitrogen) and used at dilutions 1:400 (v/v). All antibody incubations were performed in PBA containing 2 % human serum. Unless otherwise stated, all flow cytometry stainings were performed against membrane-bound DC-SIGN and Galectin-9. For intracellular DC-SIGN and Galectin-9 stainings cells were fixed in 4 % PFA (w/v) for 10 min, permeabilised by incubation for 15 min with PBA containing 0.1 % saponin (PBA-S) prior to being stained as before. For actin staining cells were fixed in 4 % PFA (w/v) for 10 min, permeabilised by incubation for 15 min with PBA containing 0.1 % saponin (PBA-S) and F-actin subsequently labelled with Alexa Fluor 647-labelled phalloidin in PBA-S for 20 min at room temperature. Cells were analysed with a FACSCalibur instrument (BD Biosciences) and results analysed using FlowJo version X software (Tree Star, Ashland, Oregon).

### Confocal microscopy

*gal9* or non-targeting siRNA –transfected moDCs were collected at day 5 and blocked with PBA + 1 % human serum prior to being incubated with primary antibody for 30 min on ice. Cells were then washed and incubated for further 30 min with the corresponding secondary antibody followed by a 60 min incubation at 12 ° C. Cells were then washed extensively, fixed in 2 % PFA and allowed to attach on poly-L-lysine-coated 12 mm glass coverslips (Electron Microscopy Sciences, Hatfield, Pennsylvania) for 30 min at room temperature. After, cells were again fixed with 2 % PFA for 20 min, washed with PBS and coverslips embedded in glass slides using Mowiol prior to be stored at 4 °C until imaging. For actin staining, cells were fixed with 4 % PFA for 20 min at 4 °C, washed twice and permeabilised by incubation with PBA containing 0.1 % saponin (PBA-S). F-actin was subsequently labelled with Alexa Fluor 647-labelled phalloidin in PBA-S for 30 min at room temperature. Confocal images were obtained in a sequential manner using a commercial Olympus FV1000 Confocal Laser Scanning Microscope with Argon (457, 488, 515 nm), and 405, 559 and 635 diode excitation lasers and a 60× oil immersion objective (UPlanSApo 60×/1.35 Oil). Images were obtained using the FW10-ASW software (Olympus, Nijmegen, Netherlands) and processed with Fiji software.

### Ventral plasma membrane staining and super resolution imaging

5×10^5^ moDCs were seeded onto 25 mm glass coverslips (Electron Microscopy Sciences, Hatfield, Pennsylvania) in phenol red free RPMI containing 10 % FBS, 1 mM ultra-glutamine, IL-4 (500 U/ml) and GM-CSF (800 U/ml) in a humidified, 5 % CO2-containing atmosphere for 48 hours. After this time, ventral plasma membranes were prepared by sonication using a Sartorius Labsonic P sonicator with cycle set at 1 and amplitude at 20 % output. First, the sonicator tip was placed in a glass beaker containing 100 ml prewarmed hypotonic PHEM buffer (6 mM PIPES, 5 mM HEPES, 0.4 mM Mg_2_SO_4_, 2 mM EGTA). Next, coverslips were held 1-2 cm below the sonicator tip at a 45 degrees angle in the hypotonic PHEM solution and cells were sonicated for approximately 1 second. Directly after sonication coverslips were transferred to a pre-warmed PBS solution containing 4 % paraformaldehyde and 0.05 % glutaraldehyde and incubated for 30 min at room temperature. For Galectin-9 and actin staining, coverslips were subsequently incubated with anti-Galectin-9 antibody (AF23045, R&D systems) for 2 h at room temperature, washed and incubated for 1 h with the secondary antibody donkey-anti-goat Alexa 568 (A11057) at 1:200 (v/v) and Alexa Fluor 488-labelled phalloidin at 1:100 (v/v) followed by DAPI staining (Sigma-Aldrich) at 1:3000 (v/v) for 10 min and at room temperature. Cells were then washed twice with PBS, one time with milliQ H_2_O, embedded onto glass slides using Mowiol (Calbiochem) and stored at 4 °C until imaging. Confocal images were obtained in a sequential manner using a commercial Zeiss LSM880 confocal scanning microscope equipped with an Airyscan Unit, 405 and 561 nm diode lasers, argon (458, 488, 514 nm) lasers and a 633 nm laser and a 63× Plan Apochromat (1.4 NA) oil immersion objective. Images were obtained using the ZEN software (Zeiss Microscopy, Breda, Netherlands) and processed with the ZEN Airyscan processing toolbox and Fiji software.

### Atomic force microscopy

6×10^5^ moDCs transfected with either Non-Targeting or *gal9* siRNA were seeded in phenol-red free RPMI supplemented as before in a 40 mm glass bottom dish (GWST-5040, WillCo, Amsterdam, Netherlands) for 48 hours. After this time, mechanical probing of cells was performed with a Catalyst BioScope (Bruker, Kalkar, Germany) atomic force microscope coupled to a confocal microscope (TCS SP5II, Leica) using the “point and shoot” feature of the Nanoscope software (Bruker). Silicon nitride cantilevers with nominal spring constants of 0.06 N/m (S-NL type D, Bruker) were used without any tip modification. The system was calibrated first in air and then in cell-free medium at 37 °C prior to each experiment by measuring the deflection sensitivity on a glass surface, which enabled determination of the cantilever spring constant using the thermal noise method (te Riet et al., 2011). Before the sample was placed, the xy movement of the sample stage was calibrated using the NanoScope software. An optical image of the cells was captured after the tip position was registered, which allowed for selection of the point (or region) of interest on the optical image. Two force − distance curves were sequentially acquired from each point selected on the membranes of stretched cells and three independent points were probed for each cell. The forward (approach) and reverse (retraction) velocities were kept constant at 1 μm/s, ramping the cantilever by 4 μm with a 3 nN threshold in a closed z loop. After baseline correction, approach curves were analysed for determination of Young’s modulus of elasticity using Sneddon’s conical indenter model (Sneddon, 1965) for which Poisson’s ratio was set as 0.5 (Lin et al., 2007) and the half angle of the indenter as 18°. Contact point-independent linearised Sneddon equation was used for fitting the approach curves (van Helvert and Friedl, 2016). The region on the approach curve through which the model was fit was determined via setting the lower and upper boundaries that corresponded to approximately 10 % and 70 % of the difference between the maximum and minimum forces exerted, respectively. Two curves per point and 2-3 points per cell were averaged to obtain Young’s modulus of elasticity per cell. Approximately 10 to 30 cells were analysed for each condition and experiment.

### Rac1-GTPase activation assay

To assess GTP-bound Rac1 levels, colorimetric G-LISA activity assay kit (BK128-S, Cytoskeleton, Denver, Colorado) was used according to manufacturer’s instructions. Day 5 moDCs were stimulated with zymosan particles (Z4250, Sigma-Aldrich) for 5, 15 or 30 min prior to being lysed in ice-cold lysis buffer, snap-frozen in liquid nitrogen and stored at -80 °C. Protein concentrations were determined using the micro BCA protein assay kit (23235, ThermoFisher scientific) and 50 μg of total protein were subsequently used for the G-LISA assay. Rac1-GTP levels were determined using the Rac1-GTP binding 96-well plates. Absorption of the wells at 490 nm was determined with an iMark microplate reader (BioRad).

### Statistical analysis

All data was processed using Excel 2013 (Microsoft) and GraphPad Prism 5 software. All Image processing was performed on ImageJ software and statistical analysis was done using Prism5. The specific statistical test used is described for each figure in the figure legend. *p* values < 0.05 were considered statistically significant.

## Acknowledgements

We thank Sjoerd van Helvert and Roel Hammink for help with atomic force microscopy. This work is supported by Grant 822.02.017 from the Netherlands Organization for Scientific Research (NWO). A. van Spriel is supported by the Netherlands Organization for Scientific Research (NWO-ALW VIDI Grant 864.11.006), the Dutch Cancer Society (KUN2014-6845), and the European Research Council (ERC CoG 724281). C. Figdor is recipient of the Netherlands Organization for Scientific Research Spinoza Prize, ERC Adv Grant PATHFINDER (269019) and KWO-award KUN2009-4402 from the Dutch Cancer Society.

## Author contributions

L.Q.C and C.G.F. and A.B.vS conceived and designed the study. L.Q.C., O.T., S.D., A.vD., S.B., B.J. and K.vdD. performed the experiments and analysed the data. T.N., and M.H. provided the galectin-9 knockout mice. L.Q.C., A.C., C.G.F. and A.B.vS. wrote the manuscript. All authors read and provided input on the manuscript.

---

**Figure supplement 1.**
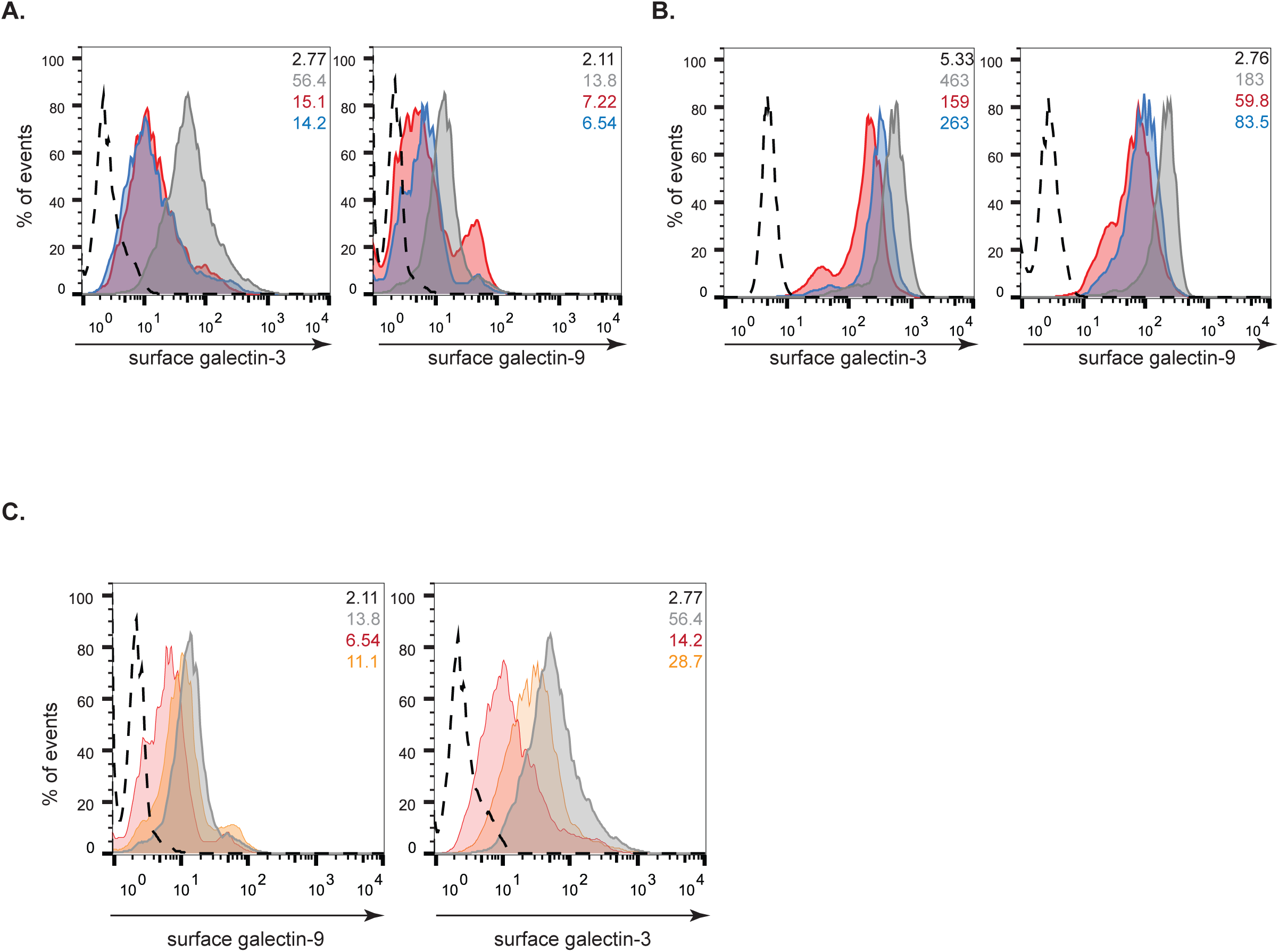
*gal9* and *gal3* siRNA do not cause non-specific effects. **A** and **B** moDCs were transfected with *gal9* siRNA and/or *gal3* siRNA or a non-targeting siRNA (NT). Galectin-9 and Galectin-3 knockdown were confirmed by flow cytometry 48 h after transfection. NT siRNA (grey area), *gal9* siRNA or *gal3* siRNA transfected moDCs (red area), *gal9* and *gal3* siRNA transfected moDCs (blue area). Black dotted line represents isotype control. Numbers in inset indicate geometrical Mean fluorescence intensity (gMFI). **C**. moDCs were transfected as above and surface levels of Galectin-9 and Galectin-3 assessed by flow cytometry. Grey area represent levels of Galectin-9 and Galectin-3 in NT siRNA transfected cells and black dotted line=isotype control. Numbers in inset indicate gMFI. In left panel: orange area shows levels of Galectin-9 in *gal3* siRNA transfected cells. Red area depicts Galectin-3 levels in *gal9* siRNA transfected cells. In right panel: organge area represents Galectin-3 levels in *gal9* siRNA transfected cells and red area shows Galectin-9 levels in *gal3* siRNA transfected cells.

**Figure supplement 2.**
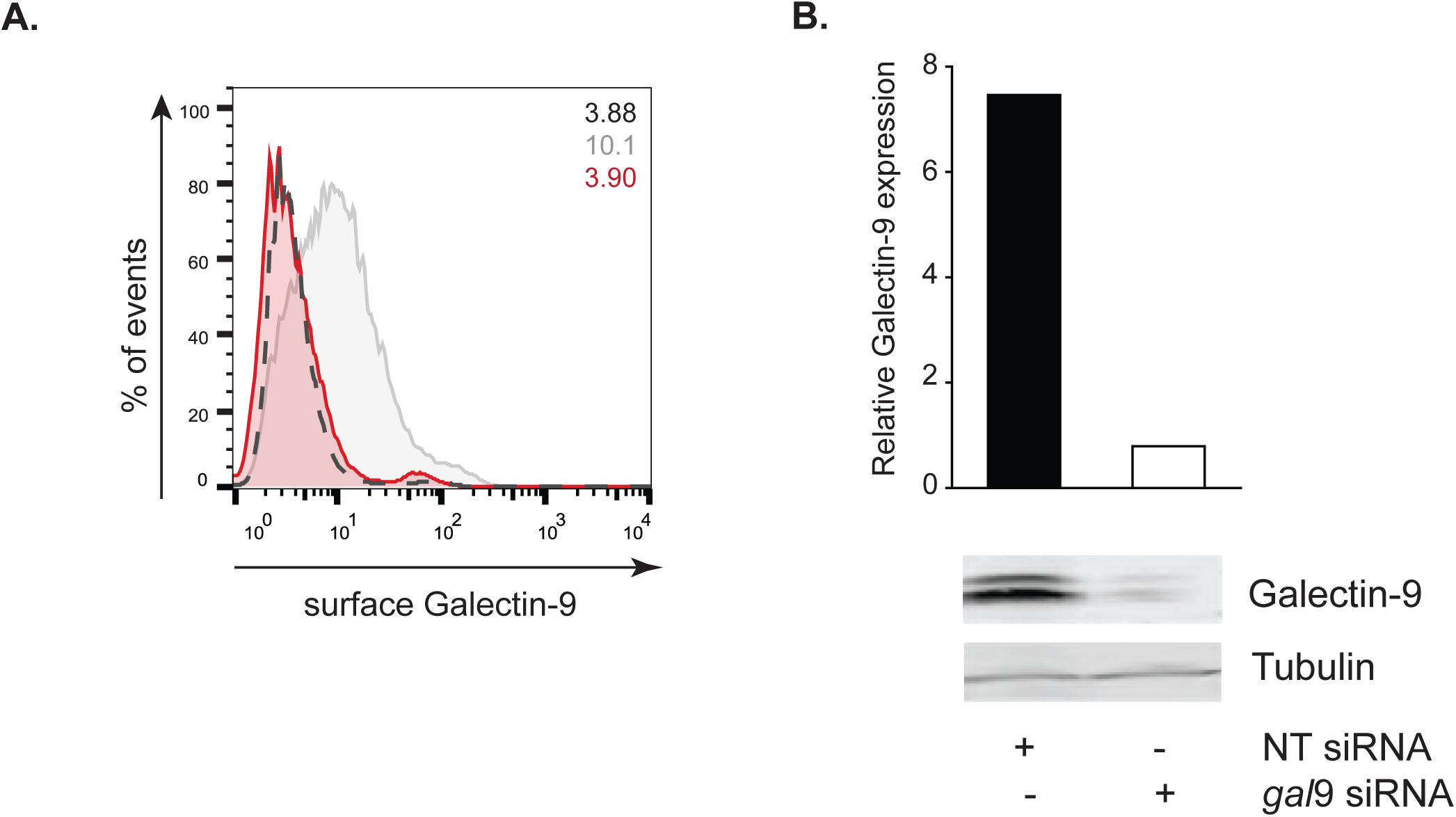
Galectin-9 is depleted in dendritic cells upon *gal9* siRNA transfection. **A** moDCs were transfected with *gal9* siRNA or a NT siRNA. Galectin-9 knockdown was confirmed by flow cytometry 48 hours after transfection. NT (grey area) or *gal9* siRNA-transfected moDCs (red area). Black dotted line represents isotype control values. Numbers in inset indicate gMFI. **B** Total lysates from NT and *gal9* siRNA transfected cells were subjected to Western Blot and Galectin-9 expression was analysed. Tubulin was used as loading control. Band intensities were quantified using ImageJ and normalised for tubulin.

**Figure supplement 3.**
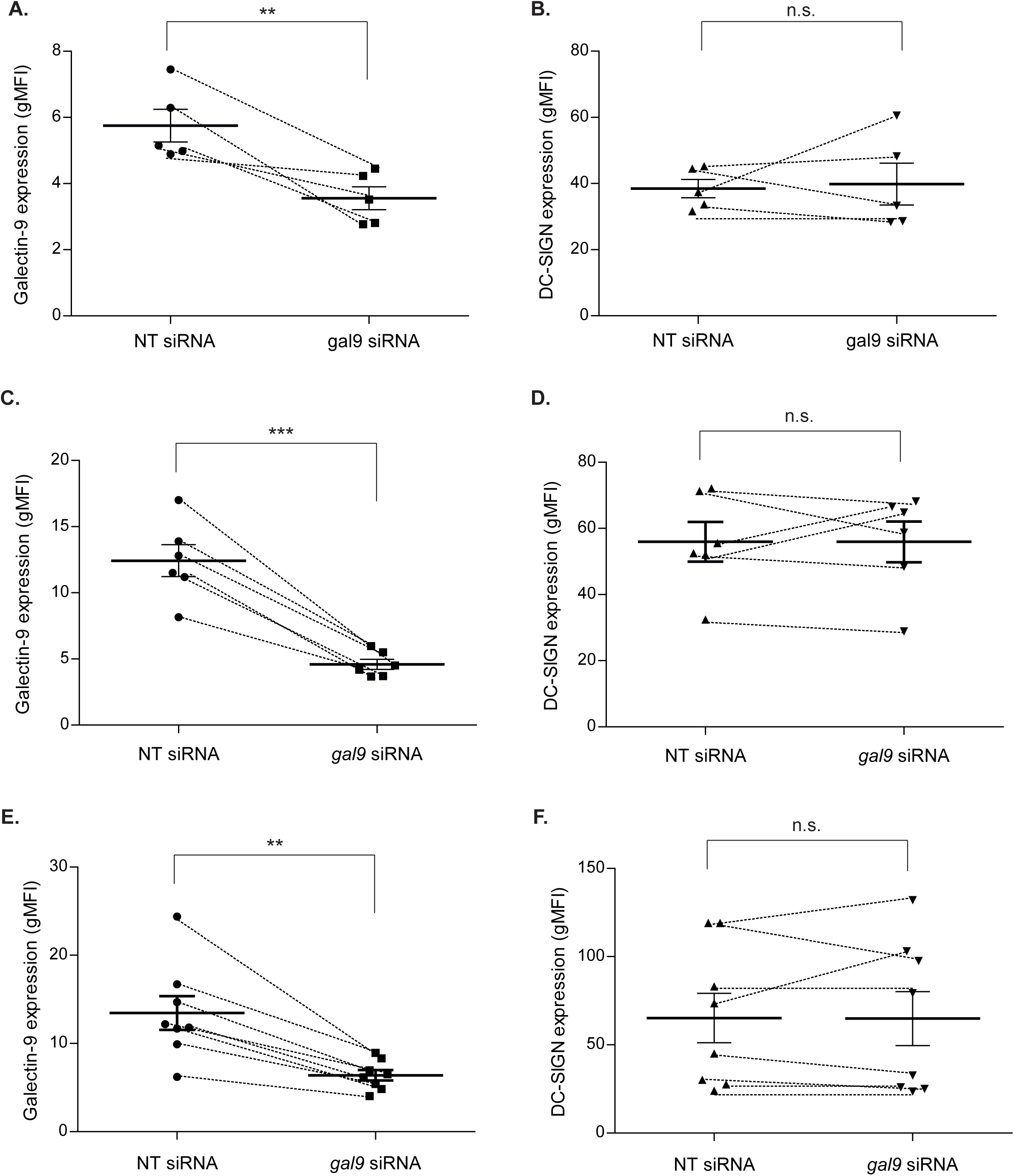
Galectin-9 knockdown does not affect cell surface DC-SIGN levels. moDCs were transfected with either NT or *gal9* siRNA and the cell surface levels of Galectin-9 and DCSIGN analysed 24 h (**A** and **B**), 48 (**C** and **D**) and 72 h (**E** and **F**) after transfection by flow cytometry. Each symbol represents one independent donor and lines connect paired NT and *gal9* siRNA-transfected moDCs. Data represents mean average expression levels ± SEM. For statistical analysis, paired students t-test was conducted between NT and *gal9* siRNA-transfected cells. n.s. p >0.05, ** p < 0.005, *** p < 0.001.

**Figure supplement 4.**
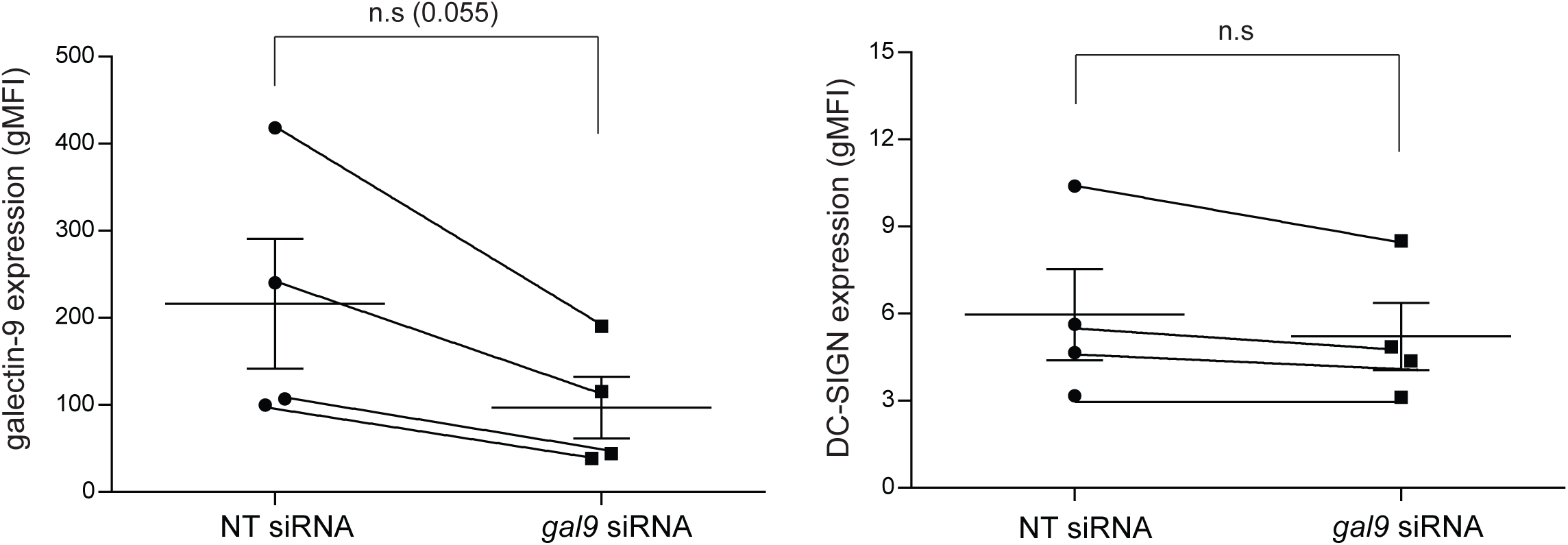
Galectin-9 knockdown does not affect intracellular DC-SIGN levels. moDCs were transfected with either NT or *gal9* siRNA and intracellular levels of Galectin-9 (**A**) and DC-SIGN (**B**) analysed 48 h after transfection by flow cytometry. Each symbol represents one independent donor and lines connect paired NT and *gal9* siRNA-transfected moDCs. Data represents mean average expression levels ± SEM. For statistical analysis, paired students t-test was conducted between NT and *gal9* siRNA-transfected cells. n.s. p >0.05.

**Figure supplement 5.**
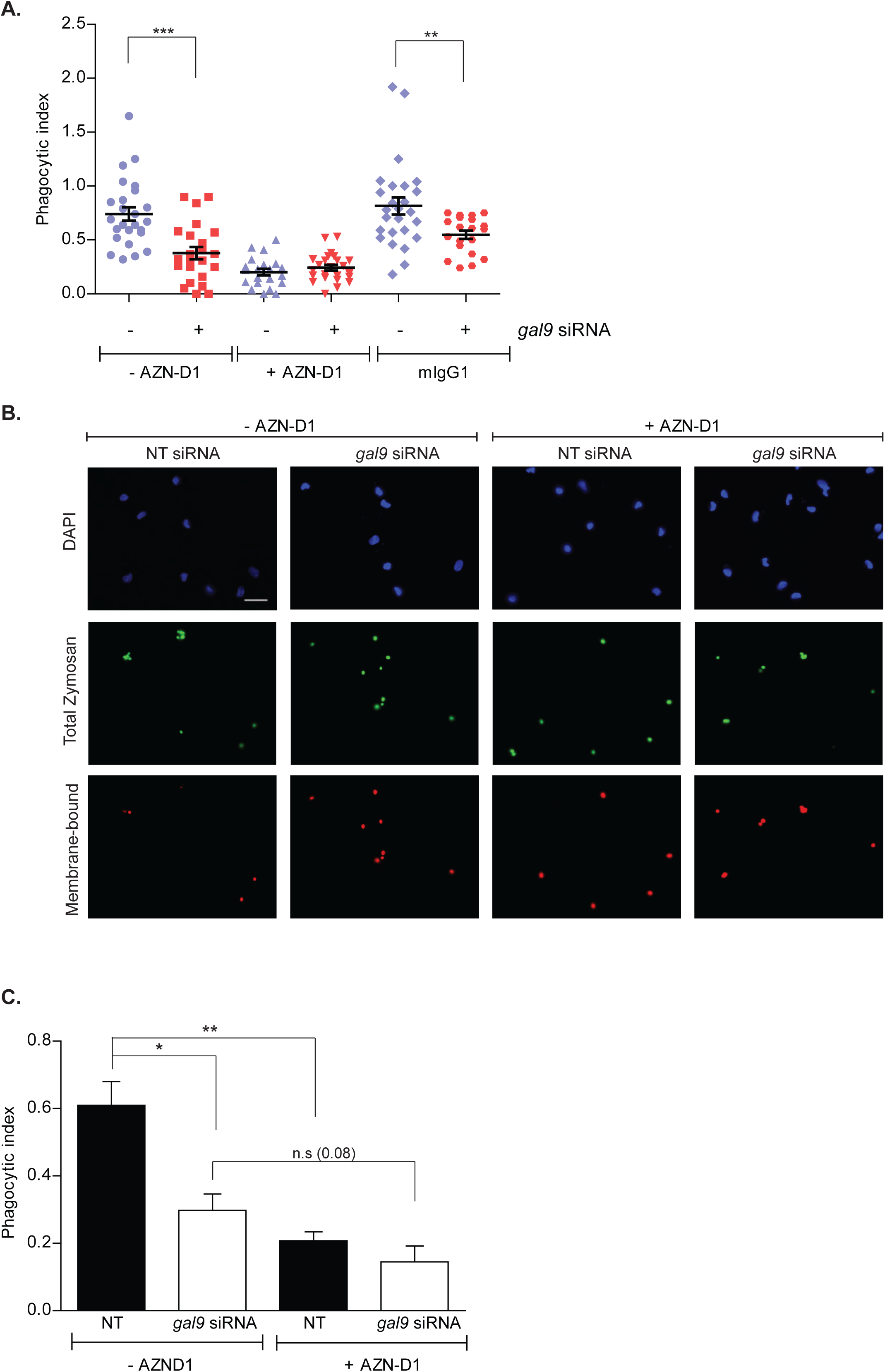
DC-SIGN is essential for particle uptake in DCs. **A** and **B** moDCs were transfected with *gal9* or a NT siRNA. Forty-eight hours later cells were incubated with AZN-D1 or isotype control (mIgG1) for 10 min prior to being challenged with zymosan for 60 min. Cells were then fixed, stained and the phagocytic index calculated. Graph and panels show representative results for one donor out of three independent experiments. Scale bar: 25 μm. **C** Quantification and statistical analysis of experiments shown in (A). Data represents mean average phagocytic index ± SEM of three independent donors. Results show the mean value ± SEM of three independent donors. Unpaired students t-test was conducted between NT and *gal9* siRNA-transfected cells. n.s p > 0.05, * p < 0.05, ** p < 0.005, *** p < 0.001.

**Figure supplement 6.**
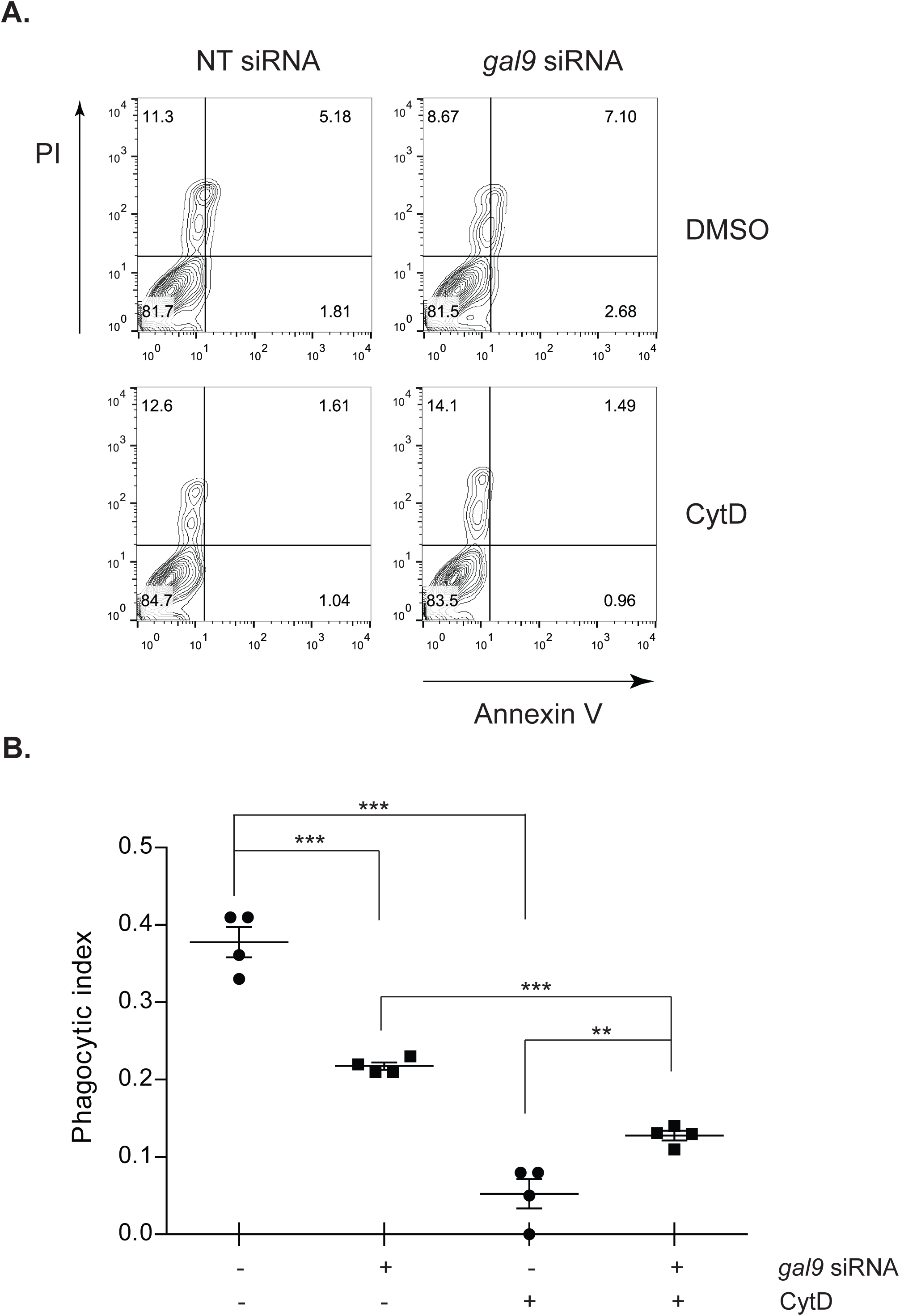
Treatment with cytochalasin D does not affect cell viability. **A** NT or *gal9* siRNA-transfected moDCs were treated with 5 μg/ml cytochalasin D (cytD) for 10 min. After this time cells were subjected to propidium iodide (PI) and Annexin V-FITC double staining for flow cytometry and percentage of apoptotic and necrotic cells calculated. One representative donor out-of-three independent experiments is shown. **B** moDCs were transfected as in (A) and pre-treated with 2.5 μg/ml cytD for 10 min prior to being challenged with zymosan for 60 min. Cells were fixed, stained and the phagocytic index calculated. Twenty frames were analysed for each condition and donor. Data represents mean average phagocytic index ± SEM for one representative donor out-of-four independent experiments. Unpaired students ttest were conducted between NT and *gal9* siRNA-transfected cells. ** p < 0.005, *** p < 0.001.

